# Bilateral symmetry of linear streptomycete chromosomes

**DOI:** 10.1101/2021.03.09.434596

**Authors:** Lis Algora-Gallardo, Jana K Schniete, David R. Mark, Iain S. Hunter, Paul R. Herron

## Abstract

Here we characterise an uncommon set of telomeres from *Streptomyces rimosus* ATCC 10970, the parental strain of a lineage of one of the earliest-discovered antibiotic-producers. Following the closure of its genome sequence, we then compared unusual telomeres from this organism with the other five classes of replicon ends found amongst streptomycetes. Closed replicons of streptomycete chromosomes were organised with respect to their phylogeny and physical orientation, which demonstrated that different telomeres were not associated with particular clades and were likely shared amongst different strains by plasmid-driven horizontal gene transfer. Furthermore, we identified a ~50 kb origin island with conserved synteny that is located at the core of all streptomycete chromosomes and forms an axis around which symmetrical chromosome inversions can take place. Despite this chromosomal bilateral symmetry, a bias in *parS* sites to the right of *oriC* is maintained across the family *Streptomycetaceae* and suggests that the formation of ParB/*parS* nucleoprotein complexes on the right replichore is a conserved feature in streptomycetes. Consequently our studies reveal novel features of linear bacterial replicons that, through their manipulation, may lead to improvements in growth and productivity of this important industrial group of bacteria.

## Introduction

Oxytetracycline (OTC) is a tetracycline analogue mass-produced under the trade name Terramycin and was one of the first antibiotics discovered. Although no longer widely used for clinical infections due to increased resistance, OTC is still used in aquaculture and a semi-synthetic derivative, doxycycline, is used against malaria and Lyme disease (Petkovic et al., 2006). The producing organism, *Streptomyces rimosus* G7, was isolated from soil near Pfizer laboratories in Connecticut, USA (Finlay et al., 1950) and deposited with NRRL as strain 2234 before being passed to ATCC. This strain, *S. rimosus* ATCC 10970, also known as G7 or R7 (Petkovic et al., 2006), was further developed for improved OTC productivity by random mutagenesis and acts as the founder of the many OTC production strains generated by strain improvement programmes (Petkovic et al., 2006). In order to form a foundation to study the development of specialised productivity, we set out to fully close the genome of *S. rimosus* G7, obtained from Pfizer. In recent years, computational tools have streamlined the elucidation of the specialised metabolic biosynthetic potential of microbial genome sequences (Chavali and Rhee, 2018). Accurate prediction of the molecules produced by these organisms is dependent on the presence of a complete genome sequence; in the case of organisms with linear chromosomes, knowledge of the telomeres of individual replicons is essential to delineate the full coding capacity of the genome.

Following the closure of the *S. rimosus* genome sequence, we noticed that the location of certain genes and sequence motifs required for effective replication and segregation of its two replicons were shared with other streptomycetes. Bidirectional DNA replication in most bacteria begins at a central origin (*oriC*) and proceeds around the chromosome until the two replication forks meet at the terminus (*ter*) of the circular molecule (Trojanowski et al., 2018). Streptomycetes possess linear chromosomes and plasmids (Lin et al., 1993) flanked by terminal inverted repeats (TIRs) that can be over 600 kb in size (Ruckert et al., 2013). Here we use the term streptomycete to refer to members the three genera that form the family *Streptomycetaceae: Streptomyces, Kitasatospora* and *Streptacidiphilus*. Archetypal telomeres are the best characterised termini of streptomycete linear replicons and contain several palindromes with differences in sequence that occur in complementary pairs of stem structures in the terminal 150 bp (Huang et al., 1998). These palindromes adopt a clover leaf structure required for priming by palindrome I in archetypal telomeres (Yang et al., 2017). However, there are also non-archetypal telomeres, such as those of SCP1; a giant linear plasmid (GLP) of *Streptomyces coelicolor*. In addition, a number of other termini have been identified that do not fit into either category from *Streptomyces griseus* 2247, *Streptomyces griseus* 13350 and two linear plasmids, pLR1 and pLR2 (Goshi et al., 2002, Ohnishi et al., 2008, Zhang et al., 2006). Whist all known telomeres contain palindromes, their sequence differs between each class (Fig. 1).

**Fig. 1.**
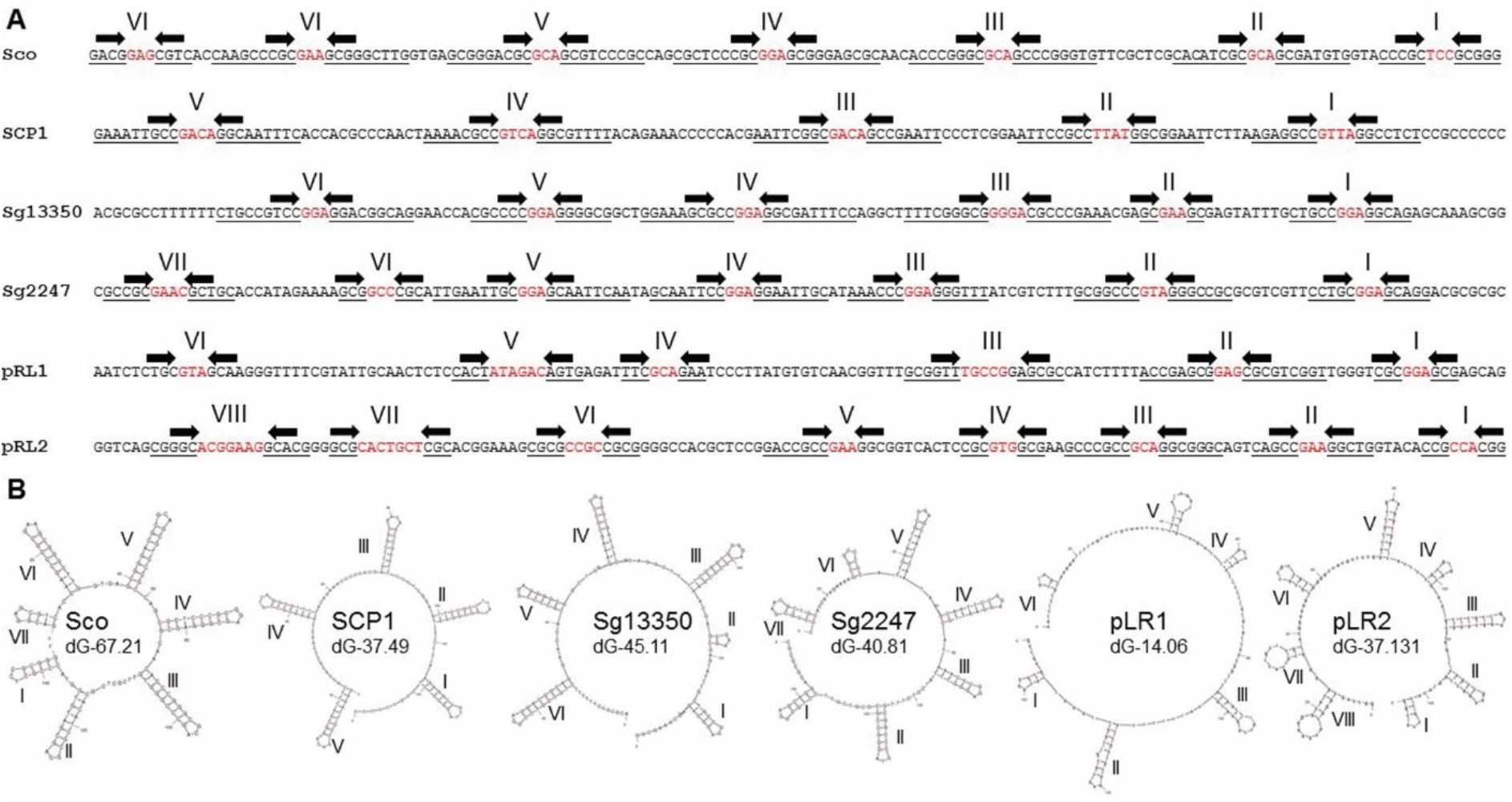
The six known classes of streptomycete telomeres. **(A)** Terminal 150bpof 3’ replicon ends of known streptomycete telomeres showing stem structures (underlined, arrows) and hairpins (red) identified using Mfold (I-VIII). Sco, archetypal end from *S. coelicolor* chromosome (Bentley et al., 2002); SCP1, non-archetypal end from *S. coelicolor* plasmid SCP1 (Huang et al., 2007); Sg13350, chromosome end from *S. griseus* 13350 chromosome (Ohnishi et al., 2008); Sg2247, chromosome end from *S. rimosus* ATCC10970 chromosome (this work); pRL1 end from *Streptomyces* sp. 44030 plasmid pRL1(Zhang et al., 2006); pRL2 end from *Streptomyces* sp. 440414 plasmid pRL2 (Zhang et al., 2006). **(B)** Mfold projections of the terminal 150 bp of the known classes of streptomycete telomeres showing stem-loop structures (I-VIII) identified using Mfold (Zuker, 2003) are displayed where the revised free energies (ΔG) were determined using Jacobson-Stockmeyer theory to assign free energies to multi-branch loops. Further details of known streptomycete telomeres are displayed in Table S2 and each individual Mfold representation in Fig. S4.

While end-patching of archetypal streptomycete telomeres is well understood and is carried out by the telomeric proteins, Tap and Tpg (Yang et al., 2017), there is still a gap in our knowledge for most other types of telomeres. Tap and Tpg are essential for the replication of linear streptomycete replicons with archetypal telomeres as their deletion leads to replicon circularization (Bao and Cohen, 2003). Tac and Tpc are required for end-patching of the non-archetypal telomeres of SCP1 (Huang et al., 2007), but there is no mechanistic information for end patching of non-archetypal or the other streptomycete telomeres.

Initially, the aim of this research was to complete the genome sequence of *S. rimosus* ATCC 10970 (Petkovic et al., 2006, Pethick et al., 2013). Key to this was to determine the telomeric sequences of the two *S. rimosus* replicons; a chromosome and GLP, first identified by pulsed field gel electrophoresis (PFGE) (Gravius et al., 1994, Pandza et al., 1997). However, when attempting a comparison with other streptomycete genomes, we were only able to identify a small number of streptomycete genome sequences flanked by known telomeres and were thus truly closed (Table S3). We then set out to analyse these closed sequences for the location of *oriC, parS* sites and genes encoding the telomeric proteins (Tap/Tpg) and partitioning proteins (ParA/ParB) (Jakimowicz et al., 2002, Kois-Ostrowska et al., 2016). In so doing, we identified shared features of streptomycete linear chromosomes that increases our understanding of how genomic architecture is conserved in this important antibiotic-producing bacterial group.

## Results and Discussion

### The closed genome of *S. rimosus* ATCC10970 consists of two linear replicons

*S. rimosus* is one of the most sequenced streptomycetes with 38 listed in NCBI at time of writing (Assembly Level: complete, 2; chromosome, 1; scaffold, 2; contig, 33), although none contain recognisable streptomycete telomeres. Analysis of 32 *S. rimosus* genome sequences shows that members of this species encode between 35 and 71 specialised metabolite biosynthetic gene clusters (Park and Andam, 2019). Many streptomycetes contain BGCs in the arms of linear chromosomes (Kinashi, 2011), whose annotation is likely complicated by the presence of multiple contigs and TIRs. Consequently we used a combination of PacBio and Illumina sequencing, in conjunction with the physical recovery and sequencing of the chromosome and plasmid ends, to generate a fully closed sequence of *S. rimosus*. The assembly details for the genome are described in the Supplementary Information.

*S. rimosus* ATCC 10970 consists of a linear chromosome and plasmid, SRP1 (Fig. S1A.), which is consistent with earlier physical studies (Gravius et al., 1994, Pandza et al., 1997). The chromosome is 9,351,267 bp in size with 11,386 bp TIRs. The GLP, SRP1, is 292,624 bp in size with TIRs of 288 bp. Together these two replicons encode 8292 CDS, 7 rRNA operons, 68 tRNAs and 3 ncRNAs. The *S. rimosus* ATCC 10970 genome is listed in NCBI under accession number CP048261. The sizes of the GLP, SRP1 was verified by PFGE and the assemblies of both replicons was verified by DraI and AseI restriction digestion followed by PFGE (Fig. S1B). The restriction patterns observed agreed with the *in silico* predictions of their size and corroborates our assembly. PFGE patterns from *S. rimosus* R7 (R7 and G7 are synonyms for ATCC 10970) were similar to those from *S. rimosus* R6 (Pandza et al., 1997) except that an additional fragment was located at one chromosome end of R7. This suggests R6 and R7 share a common ancestor and might account for the different telomeres found in *S. rimosus* R6 and R7 (Petkovic et al., 2006) and reported here.

### The replicons of *S. rimosus* ATCC 10970 are flanked by rare telomeres

In order to close the genome sequence of *S. rimosus* ATCC 10970 we employed a self-ligation PCR-sequencing method (Fan et al., 2012) to recover the telomeres of the chromosome and SRP1. Assembly of Illumina and PacBio reads produced two contigs corresponding to these replicons (see supplementary information) and allowed us to identify SmaI and PvuII as restriction enzymes that digested close to the ends of both molecules. When the *S. rimosus* genomic DNA was digested with SmaI and PvuII and the fragments were self-ligated, circular molecules were generated that corresponded to the ends of both replicons (Fig. S2). Using the primers listed in Table S1, the telomeres were recovered as PCR products and sequenced. These telomere sequences were combined with the draft sequence of both replicons to close the sequences. Here we define a closed streptomycete sequence to include the entire sequence of all replicons; many sequences listed as complete in NCBI do not contain recognizable telomeres. The chromosomal telomeres had identical sequences, whilst the left and right hand telomeres of SRP1 had a high degree of similarity (Fig. 2) to each other and to a group of telomeres first found in *S. griseus* 2247 (Goshi et al., 2002) that are distinct from archetypal and non-archetypal telomeres. Although the telomeric sequence from this strain has not been submitted to NCBI, we named this class of telomeres as Sg2247. An archetypal telomeric sequence exists in NCBI from the telomere of *S. rimosus* R6 (Huang et al., 1998) (Accession number AY043328.1) and suggests that telomeric heterogeneity exists within strains of this species. *S. rimosus* ATCC 10970 and *S. rimosus* R6 are independent isolates: the former is the original soil isolate (Finlay et al., 1950) and the latter is a soil isolate from Zagreb, Croatia (Petkovic et al., 2006).

**Fig. 2.**
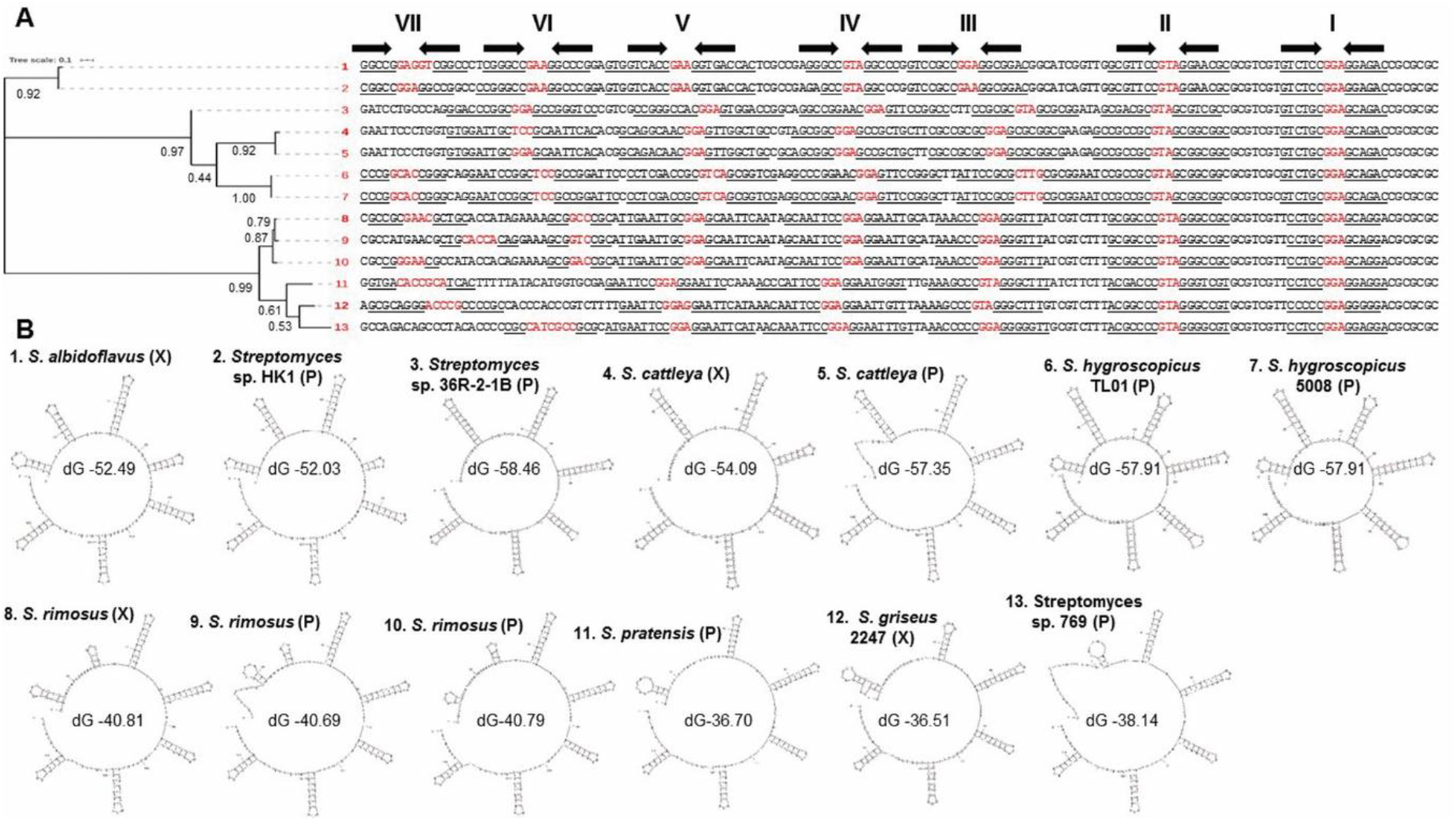
Evolutionary relationships of Sg2247-type telomeres showing stem-loop structures. **(A)** Phylogenetic tree of 3’ replicon end sequences with similarity to the telomeres of *S. rimosus* ATCC 10970, (Sg2247-type telomeres (Goshi et al., 2002)). The proportion of trees in which the associated taxa clustered together is shown next to the branches. The tree is drawn to scale, with branch lengths measured in the number of substitutions per site. **1**, *S. albidoflavus* J1074 chromosome, accession number NC_020990.1; **2**, *Streptomyces* sp. HK1 plasmid pSHK1, EU372836.1; **3**, *Streptomyces* sp. 36R-2-1B plasmid pYY8L, GU080325.1; **4**, *S. cattleya* NRRL 8057 chromosome, FQ859185.1; **5**, *S. cattleya* NRRL 8057 plasmid pSCAT FQ859184.1; **6**, *S. hygroscopicus* subsp. jinggangensis TL01 plasmid pSHJGH1 right hand end, NC_020894.1; **7**, *S. hygroscopicus* subsp. jinggangensis 5008 plasmid pSHJG1 right hand end, NC_017766.1; **8**, *S. rimosus* ATCC10970 chromosome, CP048261.1; **9**, *S. rimosus* ATCC10970 plasmid SRP1 left hand end, CP048261.2; **10**, *S. rimosus* ATCC10970 plasmid SRP1 right hand end, CP048261.2; **11**, *S. pratensis* ATCC 33331 plasmid pSFLAO1, CP002476.1; **12**, *S. griseus* 2247 chromosome; **13**, *Streptomyces* sp. 769 plasmid pSGZL, CP003988.1. **(B)** The terminal 150bp of 3’ replicon ends of Sg2247-type telomeres showing stem structures and hairpins identified using Mfold (Zuker, 2003) are displayed. Revised free energies (ΔG, kJ/mol) were determined using Jacobson-Stockmeyer theory to assign free energies to multi-branch loops. Those telomeres located on chromosomal (X) and plasmid (P) replicons are designated in parentheses. Further details of known Sg2247 telomeres are displayed in Table S2 and each individual Mfold representation in Fig S3.

We next set out to determine the incidence of Sg2247 type telomeres across the streptomycetes. We used the terminal 150 bp of the *S. rimosus* chromosome telomere to search NCBI using BLASTn. This generated over 50 hits, with a query coverage and identity of at least 19% and 76% respectively. To exclude pseudo-telomeres (Huang et al., 2003, Huang et al., 2007) and detect true telomeric sequences, we only included sequences that were found at the end of a replicon from complete genomes or where the telomere sequence had been physically isolated and sequenced. 11 replicons contained telomeres similar to those of *S. rimosus* (Sg2247 type); two were at both chromosome ends (*S. rimosus* ATCC 10970 and *S. albidoflavus* J1074 (Zaburannyi et al., 2014)), whilst *S. cattleya* NRRL8057 (Barbe et al., 2011) possesses a Sg2247 telomere at one end and no recognizable telomere at the other. In addition, Sg2247 type telomeres were found at both termini of three plasmids and at one terminus of a further four plasmids. Interestingly two plasmids (pSHJG1 from *S. hygroscopicus* subsp. jinggangensis 5008 (Wu et al., 2012) and pSHJGH1from *S. hygroscopicus* subsp. jinggangensis TL01) contained a Sg2247-type telomere at one terminus and an archetypal telomere at the other.

A ClustalW alignment of 13 Sg2247 telomeres was used to determine the evolutionary relationships within this class of telomeres (Fig. 2A). Sg2247-type telomeres form similar stem-loop structures to those found in archetypal and non-archetypal telomeres (Huang et al., 2007), with up to eight stem-loop structures predicted within the terminal 150bp and highest similarity at their 3’ ends where terminal stem-loop structures are highly conserved (Fig. 2, individual Mfold projections are shown in Fig. S3). The most common sequence found at the hairpins was GGA, GAA and GTA, similar to that found at the hairpins of archetypal telomeres (Huang et al., 2007). Revised free energies (ΔG) of the stem-loop structures predicted by Mfold (Zuker, 2003) varied between −36.51 and −58.46 kJ/mol. Sg2247 telomeres fell into two groups that reflected their phylogenetic relationship (Fig. 2B): those with a relatively low ΔG (strains 1-7) and those with a significantly higher ΔG (strains 8-13, according to a Students t-test (p< 6.47×10^−7^).) Strains with the lowest ΔG (1-7) contained longer hairpins distal to the 3’ replicon end, although it is unclear whether this has any functional significance.

### Six classes of telomeres are found in the linear replicons of streptomycetes

Archetypal telomeres are found in many streptomycete replicons, such as the chromosome end of *S. coelicolor* M145 (Bentley et al., 2002). Non-archetypal telomeres were first identified in SCP1, the GLP of *S. coelicolor* A3(2) (Huang et al., 2007) and another class of telomeres was identified in the *S. griseus* 13350 chromosome (Ohnishi et al., 2008), here termed Sg13350-type. Finally, two types of telomeres were found at the ends of two linear plasmids, pRL1 and pRL2 from streptomycete soil isolates (Zhang et al., 2006). Using the same approach with which we identified Sg2247-type telomeres, we also compiled a list of telomeres from all six classes that were found at the end of closed replicons or had been recovered physically and sequenced by Sanger sequencing (Fan et al., 2012). The predicted streptomycete telomeres of all six classes are listed in Table S2. Archetypal telomeres are the most frequent and are found in both chromosomes and linear plasmids, whilst three non-archetypal telomeres were identified (SCP1 (Huang et al., 2007), pSCO2 (Ruckert et al., 2013) and pFRL3 (Chen et al., 2013, unpublished data, Accession number, KF602048.1), all from linear plasmids. The chromosomal telomere of *S. griseus* 13350 (Ohnishi et al., 2008) remains the sole member of the Sg13350 class as do the telomeres of pRL1 and pRL2 of their eponymous classes (Zhang et al., 2006).

Folding analysis of the six classes of telomeres indicated that all could form stem-loop structures (Fig. 1 and individual Mfold projections in Fig. S4). Although there is little sequence similarity in the stems, the hairpins of the stem-loop structures across all six classes of telomeres are usually 3 or 4 bp in length, with GAA, GGA and GTA the most common hairpin. With the exception of Tap/Tpg (Yang et al., 2017) and Tac/Tpc (Huang et al., 2007), which prime end-patching of archetypal and non-archetypal telomeres respectively, no other telomeric proteins have been characterised. Although a putative Tap was described in *S. albidoflavus* J1074 (Zaburannyi et al., 2014), we were unable to detect any similarity to Tap, Tpg, Tac or Tpc in genome sequences with Sg2247-type telomeres. This was also true in genomes with the Sg13350 and pLR1 types of telomeres, although pRL2 (Zhang et al., 2006) encodes two candidate telomeric proteins: one displaying similarity to Tap and a helicase of *Thiobacillus* sp., whilst the other protein resembles Tpg and part of the adenovirus telomere terminal protein (Zhang et al., 2006). Taken together this indicates that, although streptomycete telomeres of all types bear some resemblance to each other in terms of the capacity of their 3’ overhangs to fold back on themselves, there are likely to be undiscovered systems able to catalyse end-patching.

Analysis of complete streptomycete genome sequences recovered few replicons that were flanked by a member of these six telomere classes. Indeed, on the basis of our strict criteria (complete sequence of all replicons and all replicons flanked by known telomeres), we could only confirm 20 closed genome sequences from streptomycetes. Many complete genomes within NCBI do not encode Tap and Tpg, so it is possible that other classes of telomeres await discovery. In a small scale study using PCR, 8 of 17 newly-detected streptomycete linear plasmids lacked typical telomeric *tap* and *tpg* sequences (Zhang et al., 2006), whilst two novel telomeres (pRL1 and pRL2) were discovered in these eight strains and one archetypal telomere from the other strains suggesting that novel streptomycete telomeres are common. If the ends of these replicons analysed during this study are representative of those of the entire family, this analysis suggests that archetypal telomeres are the most common and phylogenetically widespread as they are found in two genera (*Streptomyces* and *Kitasatospora*) of the family *Streptomycetaceae*. It seems that Sg2247–types are less abundant and form a minority of the ends of both chromosomes and plasmids, whilst the other classes of telomeres are likely to be rare.

### Streptomycete telomeres are not distributed according to the evolutionary relatedness of different strains

To determine if the six different classes of streptomycete telomeres were associated with different phylogenetic clades within the family *Streptomycetaceae*, we first compiled a list of closed streptomycete sequences. Our definition of a closed genome sequence was that all replicons of that organism must be completely sequenced and that each linear replicon was flanked at both ends by one of the six classes of telomeres described previously. At time of analysis (April 2020) there were 223 streptomycete genomes that were listed as complete in NCBI (one from the genus *Streptacidiphilus*, four from *Kitasatospora* and 218 from *Streptomyces*). After inspecting these genomes for the presence of telomeres located at both ends of all linear replicons, we identified 20 strains that met these criteria (Table S3). The replicons of other genome sequences designated as complete might be flanked by so far undiscovered telomeres or may lack identified telomeres due to the assembly difficulties of using paired-end reads to sequence linear molecules and the bioinformatic complexity introduced by the presence of TIRs at the end of replicons. We next organised all linear replicons carried by these 20 strains so that they were placed in the same direction on the basis of the *oriC* region of *S. coelicolor* (Bentley et al., 2002, Huang et al., 2013) (Fig. 3). Replicons were orientated so that *dnaN* and *dnaA* were transcribed with the bottom strand as coding strand and *parA* and *parB* transcribed with the top strand as the coding strand. Chromosomes were organized in this orientation to comply with the established orientation for *S. coelicolor* M145, which has *SCO0001* at the left and *SCO7846* on the right (Bentley et al., 2002). Subsequently, we organised the 20 closed strains phylogenetically using OrthoANI (Lee et al., 2016) and AutoMLST (Alanjary et al., 2019) with *K. setae* KM-6054 as an outgroup by generating a maximum likelihood tree and determining the location of strains with different classes of telomeres on this tree (Fig. S5). The high-resolution species tree shows that, although archetypal telomeres are the most common of the chromosome ends, two strains (*S. albidoflavus* J1074 and *S. rimosus* ATCC10970) carry chromosomes with Sg2247-type telomeres. The phylogenetic separation of these strains suggests that telomeric exchange through horizontal gene transfer has occurred. The telomeres of *S. griseus* NBRC13350 were physically recovered (Ohnishi et al., 2008) and, at time of writing, no other telomeres with similarity to those of this strain have been identified. Despite this, the closely related strains *S. annulatus* ATCC11523 and *S. globisporus* C-1027 (Fig. S5), possess archetypal telomeres. *S. rimosus* ATCC10970 (Sg2247), *S. malysiensis* DSM4137 (archetypal) and *S. binchenggensis* BCW-1 (archetypal) are also phylogenetically related, but possess telomeres from different classes. The only closed streptomycete genome from outside the genus *Streptomyces*, is from *K. setae* KM6054 with archetypal telomeres; there are no characterised telomeric sequences available for *Streptacidiphilus*. Both *S. coelicolor* M145 and *S. collinus* Tu365 possess chromosomes with archetypal telomeres, but plasmids with non-archetypal telomeres (Fig. 3). Intriguingly, the two closely related strains of *S. hygroscopicus* subsp. jinggangensis (Wu et al., 2012) possess a chromosome with archetypal telomeres, but a hybrid GLP with one archetypal and one Sg2247-type telomere. Hybrid replicons are not unknown in streptomycetes as demonstrated by the construction of *S. coelicolor* 2106 that carries a 1.85 Mb GLP (derived from SCP1), in addition to a 7.2 Mb linear hybrid chromosome with ends from the chromosome and GLP and thus this strain possesses replicons with both archetypal and non-archetypal telomeres (Yamasaki and Kinashi, 2004).

**Fig. 3.**
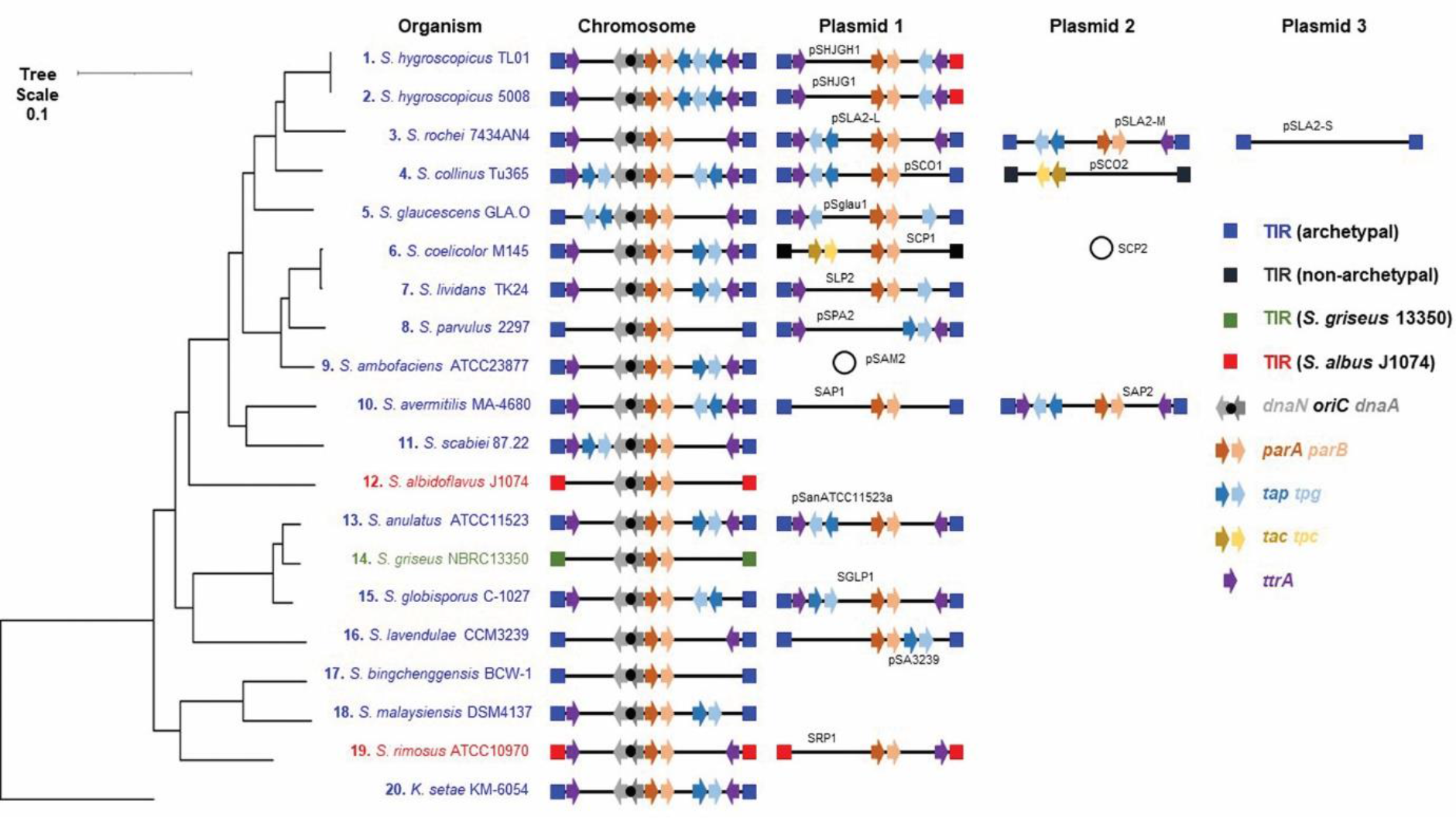
Relative position of genes encoding telomeric and partitioning proteins with respect to *oriC* and telomere class in the family *Streptomycetaceae*. All replicons carried by the 20 organisms with closed genomes (both chromosomes and plasmids) were investigated for the presence of genes encoding the archetypal and non-archetypal telomeric proteins, Tap/Tpg and Tac/Tpc respectively. In addition, genes encoding TtrA, DnaN, DnaA, ParA, ParB are plotted (Table S3). The four of the six different classes of streptomycete telomeres are also displayed (archetypal, blue; non-archetypal, black; Sg2247, red and Sg13350, green). The locations of these genes and telomeres were plotted in relation to the whole genome MLSA analysis described in Fig. S5.

We used this collection of closed genomes to investigate the TIRs by alignment of both ends of all replicons. The 20 chromosomes had a mean size of 8,846,536.25 bp (range 6,841,649-11,936,683 bp) and were flanked with 37,990 (range 13-237,155 bp) perfect inverted repeats and 77,092 (range 138-631,365 bp) imperfect inverted repeats, whilst the 16 GLPs spread over 12 strains were on average 188,460 bp in size (range 17,526-617,085 bp) with TIRs with 5,775 bp mean perfect inverted repeats and 6,552 bp mean imperfect inverted repeats. On average, a perfect TIR constituted 0.34% (0.87% imperfect) of a streptomycete chromosome and 3.06% (3.48% imperfect) of a GLP.

Whilst all chromosomes encode *dnaN, dnaA, parA* and *parB*, not all plasmids carry *parAB* (pSLA2-S, *S. rochei*; pSCO2, *S. collinus* and pSPA2, *S. parvulus*); presumably these plasmids do not require these proteins for partitioning, or their functions are provided *in trans* by the chromosomal copies of these genes. We also mapped genes encoding known telomeric proteins (*tap*/*tpg, tac*/*tpc* and *ttrA* (Fig. 3)). Non-archetypal telomeres flank two replicons, the GLPs SCP1 from *S. coelicolor* and pSSCO2 from *S. collinus*. Neither of these replicons encode Tap, Tpg or TtrA, whilst both carry *tac* and *tpc*, encoding the non-archetypal telomeric proteins (Huang et al., 2007). Consistent with previous analysis of *S. albidoflavus* J1074 (Zaburannyi et al., 2014), we were unable to locate either *tap* or *tpg* in the genomes of the two strains with Sg2247-type telomeres (*S. albidoflavus* J1074 or *S. rimosus*) or *S. griseus* 13350 (Ohnishi et al., 2008). This suggests that their end-patching is carried out by a different mechanism to that of archetypal or non-archetypal telomeres and the identity of the proteins that catalyse this process remains elusive at the present time. The two closely related strains of *S. hygroscopicus* subsp. jinggangensis (Wu et al., 2012) contain plasmids with one archetypal end and one Sg2247-type telomere. These two strains carry two copies of *tap* on the chromosome and a copy of *tpg* on both the chromosome and plasmid. All strains with archetypal telomeres encode Tap and Tpg on at least one replicon, with the exception of *S. bingchenggensis* BCW-1(Tsai et al., 2011); it is unclear how this strain primes replication at the chromosome ends. Some strains (*S. rochei* 7434AN4 (Nindita et al., 2019, Nindita et al., 2015), *S. parvulus* 2297, *S. lavendulae* CCM3239) do not carry *tap* and *tpg* on the chromosome, but do so on at least one GLP suggesting that these plasmids are required to encode the telomeric proteins for chromosomal end-patching. Conversely, several GLPs do not encode telomeric proteins: SLP2 from *S. lividans* (Tap), pSgalu1 from *S. glaucescens* GLA.0 (Tap), pSLA2-S from *S. rochei* 7434AN4 (Tap and Tpg) and SAP1 from *S. avermitilis* (Tap and Tpg) and suggest that end-patching is carried out by proteins encoded *in trans* or by other unknown proteins. Nevertheless, it seems that there is some cross-talk of the telomeric proteins and telomeres of different replicons contained within an individual strain.

Finally *ttrA*, when present on a replicon, is often found towards the ends of replicons that are flanked by archetypal telomeres and is transcribed in an inward direction suggesting that its orientation with respect to the telomeres is important for its function or transcriptional regulation. Exceptions to this are the *S. parvulus* 2297 and *S. bingchenggensis* BCW-1 chromosomes, although the former strain carries a plasmid (pSPA2) that perhaps provides TtrA functionality *in trans*. This gene is less common at the ends of replicons with non-archetypal, Sg13350-or the Sg2247-type replicon of *S. albidofvaus* J1074. Despite this, S*. rimosus* encodes a copy of *ttrA* at each chromosome termini and one copy at the right end of SRP1; unusually the latter gene is transcribed in an outward direction. The function of TtrA, with similarity to DEAD-box helicases, was hinted at through its requirement for conjugation of SLP2 in *S. lividans* (Huang et al., 2003) where both copies of *ttrA* were necessary for SLP2mediated conjugation. Consequently, the location and importance of *ttrA* for plasmid - mediated conjugation and presence of *ttrA* at both ends of most streptomycete replicons suggests that telomere exchange may have arisen through recombination of recipient chromosomes with incoming GLPs.

The covalent association of the telomeric proteins with the 5’ end of streptomycete linear replicons results in their replication being semi-conservative with respect to DNA, but conservative with respect to telomeric proteins (Tsai et al., 2011). The circular genetic maps of streptomycetes provides circumstantial evidence for association between chromosome ends and is supported by fluorescence *in situ* hybridisation studies in *S. coelicolor* (Yang and Losick, 2001). The telomeric proteins display both an intramolecular (Tpg-Tpg) and an intermolecular (Tpg-Tpc) physical association between different telomeric proteins that was demonstrated by chemical cross-linking (Tsai et al., 2011). This presents a problem for replicon segregation through the formation of replicon pseudo-dimers brought about by the *in trans* association of the telomeric proteins (Fig. 4). In a mechanism first proposed in 2011, these pseudo-dimers can only be resolved through disassociation and association of the telomeric proteins with those of the other daughter replicon or the occurrence of recombination between the two replicons in order to resolve the pseudo-dimer (Tsai et al., 2011). The appearance of replicons with different telomeres reported here might then be brought about by resolution of pseudo-dimers by recombination.

**Fig. 4.**
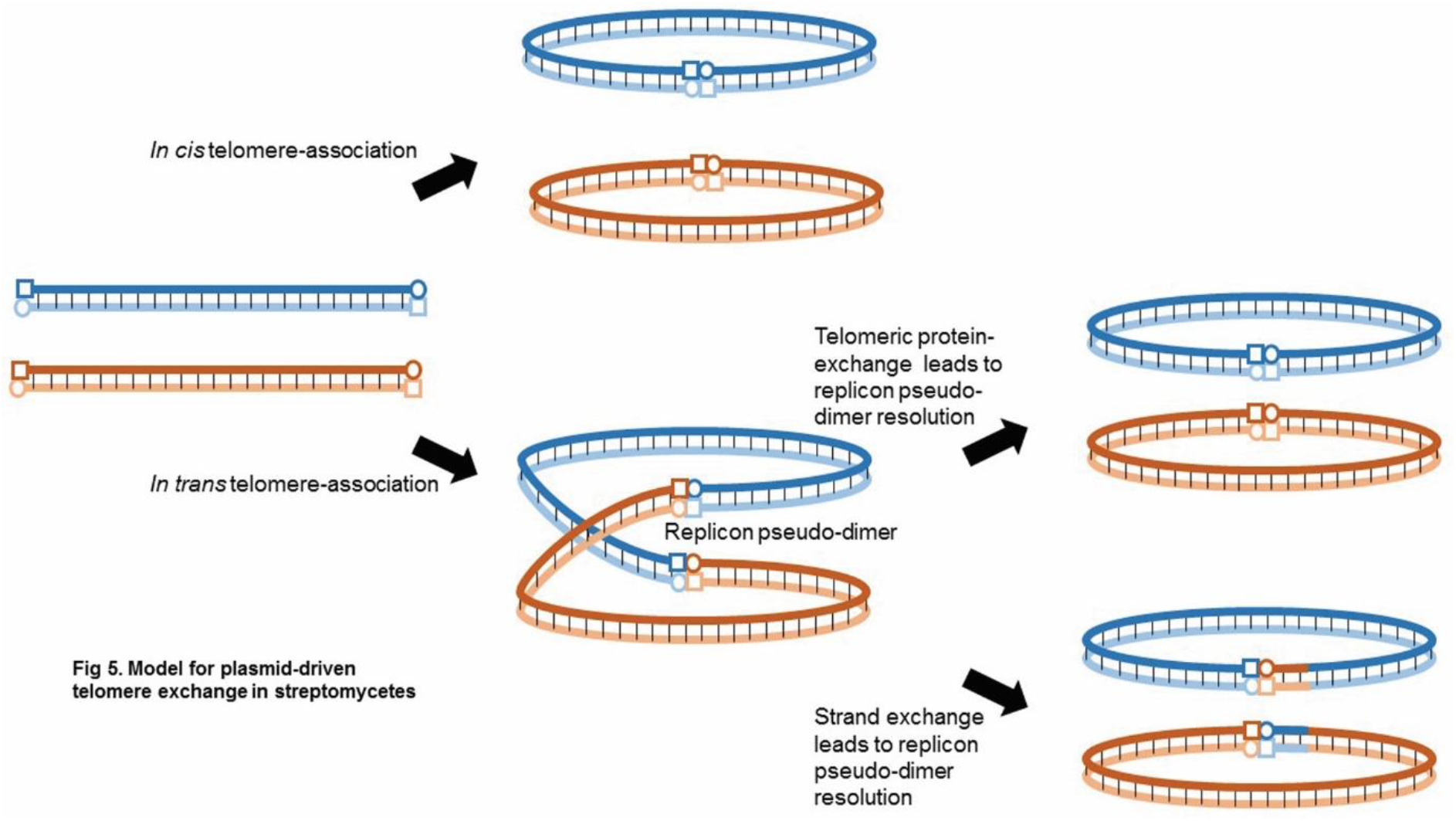
Model for telomere exchange in streptomycetes. Tpg (square) and Tap (circle) are associated with the 5’ and 3’ ends of two replicons with archeytpal telomeres (blue, brown). The telomeric proteins can either associate through an *in cis* or an *in trans* interaction. In the former case both replicons remain physically separated, however the latter results in the formation of a replicon pseudo-dimer brought about by the association of the telomeric proteins. These pseudo-dimers can only be resolved through disassociation and association of the telomeric proteins with those of the other replicon or the occurrence of recombination between the two replicons in order to resolve the pseudo-dimer. Thus, the appearance of replicons with different telomeres might then be brought about by such a genetic exchange. This model is adapted from one first proposed in 2011 (Tsai et al., 2011).

Genome plasticity and evolution of streptomycetes is generated through horizontal gene transfer in the environment and actioned by non-homologous end joining and integration of genetic material at the healing site (Hoff et al., 2018). Following a recent investigation of natural streptomycetes, it was revealed that closely related strains showed a large diversity of TIRs that reflected telomere exchanges between the chromosome and GLPs and suggests that streptomycete telomeres display extensive allelic exchange brought about by horizontal gene transfer (Tidjani et al., 2020). Strains that have undergone chromosomal changes produce diversified secondary metabolites and secrete more antibiotics (Zhang et al., 2020), so perhaps it is the capacity for genetic exchange brought about by linear replicons that contributes to the metabolic diversity and bioactivity of this bacterial group.

As a result, this analysis of closed streptomycete genomes demonstrates the diversity of telomeres and their lack of correlation with whole genome based phylogeny. Taken together with the consistency of the location of *ttrA* at both chromosomal and plasmids ends, coupled with the ability of GLPs to form hybrid ends (*S. hygroscopicus* subsp. jinggangensis), this analysis suggests that telomeric exchange between streptomycetes is a common occurrence.

### A conserved origin island lies at the centre of streptomycete chromosomes

When the 20 closed streptomycete chromosomes were orientated with *dnaA* and *dnaN* transcribed in a right to left direction, we noticed that *parA* and *parB* were transcribed in the opposite direction (Fig. 3). This suggested that the region containing *oriC* possessed conserved synteny across the analysed sequences. For this reason, we decided to investigate the extent of this synteny by carrying out a progressive alignment of the 20 closed chromosomes using Mauve (Darling et al., 2010). This analysis showed a ~50 Kb locally collinear block (LCB), containing *oriC*, that lay in the core region of all chromosomes (brown arrow, Fig. S6), whilst other LCBs (green, purple arrows, Fig. S6) in the flanking regions displayed bilateral symmetry around *oriC*. For example, the green and purple LCBs occur in a similar orientation flanking *oriC* in closely related strains. In general, the green and purple LCBs are located on opposite replichores, with the exception of the related strains *S. bingchenggensis* BCW-1 *S. malaysiensis* DSM4137 where both the green and purple LCBs are located on the same replichore. This suggests that recombination cannot occur within this core (brown) LCB and that, at the core of streptomycete chromosomes, there is a conserved region of ~50Kb that we have designated the origin island.

We next investigated chromosome synteny in more detail through the generation of dotplots using Nucmer (Marcais et al., 2018) where the chromosomal synteny of all strains was compared with each other and organized according to the MLSA tree for the 20 closed chromosomes displayed in Fig. S5. Dotplots for all 20 chromosomes are shown in Fig. S7 and for selected phylogenetic nodes in Fig. 5 that best illustrated chromosome synteny between related strains. Comparison of dotplots between sequences from related strains showed synteny across the entire length of the chromosomes, such as with the *S. hygroscopicus* node (Fig. 5A). Synteny is reduced at the termini in comparison with the core region of the chromosome and likely reflects the genetic compartmentalisation of streptomycete chromosomes that is consistent with the demonstration of a link between chromosome folding and gene expression in *S. ambofaciens* (Lioy et al., 2021). However, comparison of strains located at other phylogenetic nodes (Fig. 5B, C, D) show that one or more major chromosome inversions had taken place during the evolutionary divergence of *S. coelicolor*/*S. lividans* and *S. parvulus/S. amobofaciens* (Fig. 5B). Similar inversions were also identified at the *S. griseus* (Fig. 5C) and *S. rimosus* nodes (Fig. 5D). Interestingly, although the *S. griseus* 13350 chromosome carries the unique Sg13350 telomere class, its high average nucleotide identity with both *S. anulatus* and *S. globisporus*, which both possess archetypal telomeres (Fig. S5), suggests that telomere exchange was a relatively recent evolutionary event. The obvious bilateral symmetry between related chromosomes suggests that the recombination events leading to these rearrangements take place between the two replichores on either side of the origin island. It may be, therefore, that the conserved synteny of the origin island represents an irreducible axis around which recombination can take place, whilst not being subject to recombination itself. It also suggests that maintenance of an intact origin island is necessary for successful chromosome replication, segregation or maintenance of proper genomic topology for control of gene expression in different chromosomal regions (Lioy et al., 2021).

**Fig. 5.**
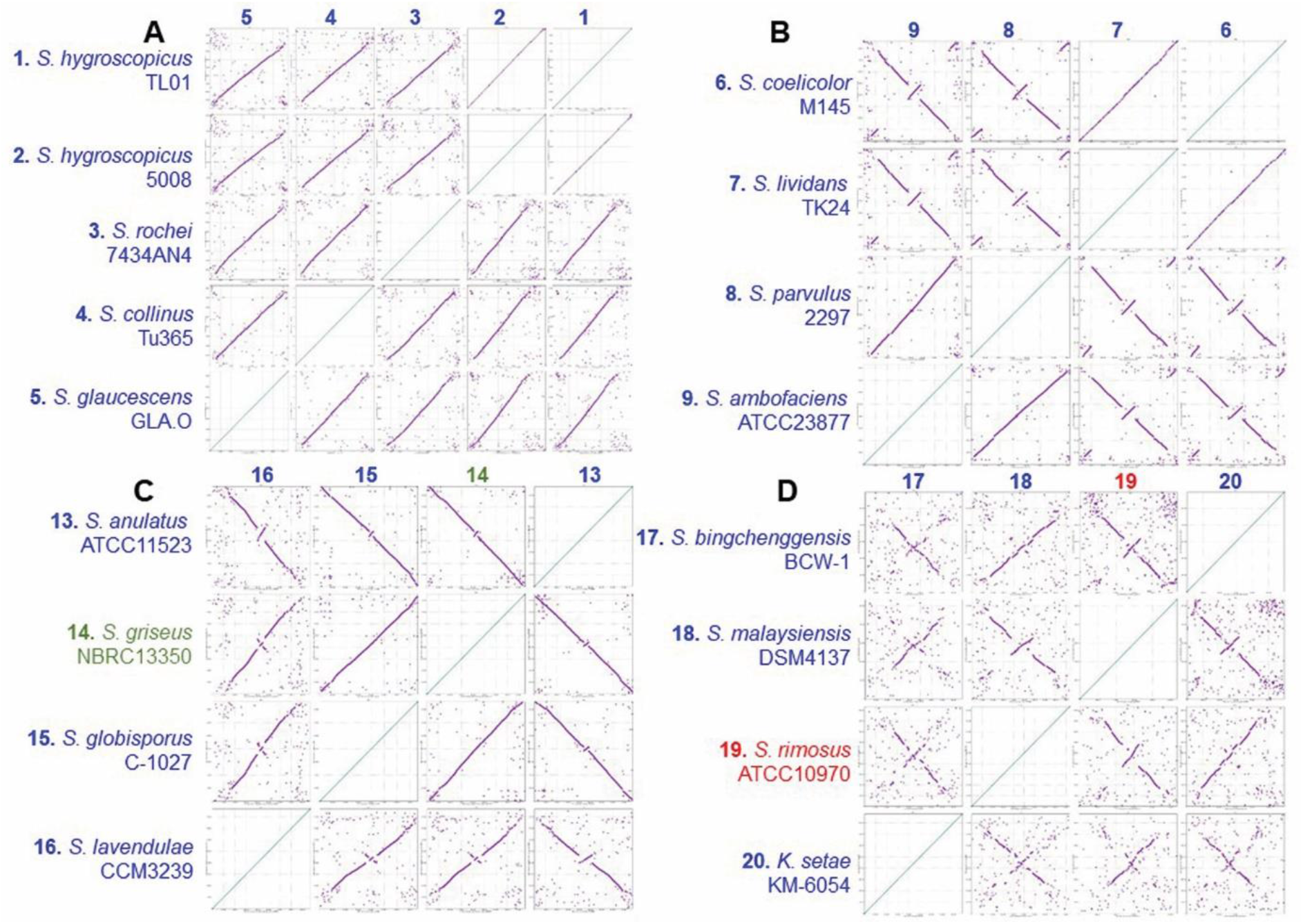
Related strains display both syntentous and asynetenous chromsomes. Each of the 20 closed streptomycete chromosomes, organized, so that the *oriC* regions were syntenous, were submitted to Nucmer and dot plots generated (Marcais et al., 2018). A comparison of all sequences is displayed in Fig. S7. Dot plots of strains around selected phylogenetic nodes (Fig. S5) are displayed. **A**; 1, *S. hygroscopicus* subsp. *jinggangensis* TL01; 2, *S. hygroscopicus* subsp. *jinggangensis* 5008; 3, *S. rochei* 7434AN4; 4, *S. collinus* Tu 365; 5, *S. glaucescens* GLA.O. **B**; 6, *S. coelicolor* M145; 7, *S. lividans* TK24; 8, *S. parvulus* 2297; 9, *S. ambofaciens* ATCC 23877. **C**; 13, *S. anulatus* ATCC 11523; 14, *S. griseus* subsp. *griseus* NBRC 13350; 15, *S. globisporus* C-1027; 16, *S. lavendulae* subsp. *lavendulae* CCM3239. **D**; 17, *S. bingchenggensis* BCW-1; 18, *S. malaysiensis* DSM4137; 19, *S. rimosus* ATCCC10970; 20, *K. setae* KM-6054.

In *S. coelicolor* this origin island contains the region corresponding to *SCO3872-SCO3911* (Fig. 6) and includes *gyrA, gyrB, recF, dnaN, oriC, dnaA, parA, parB, ssbA and dnaB*. Across all 20 strains the mean origin island size (Table S4) was 51,055 bp and constituted 0.59 % of the chromosome and its location was on average 48.49 % (range 41.25 %-53.25 %) from the left-hand end of the replicon. Multiple genome alignments of streptomycete chromosomes show that they contain highly conserved core regions and variable sub-telomeric regions (Bu et al., 2020) and it has previously been reported that the central part of streptomycete chromosome are highly syntenic (Choulet et al., 2006). Whilst symmetrical rearrangements of streptomycete chromosomes have been reported before (Zaburannyi et al., 2014, Wu et al., 2012), this is the first time that they have been related to a central axis. This suggests that the integrity of the origin island is important for maintenance of correct chromosome structure and function as when chromosome inversions take place, they do so in a symmetrical manner (between replichores) rather than in an asymmetrical manner (within a replichore).

**Fig. 6.**
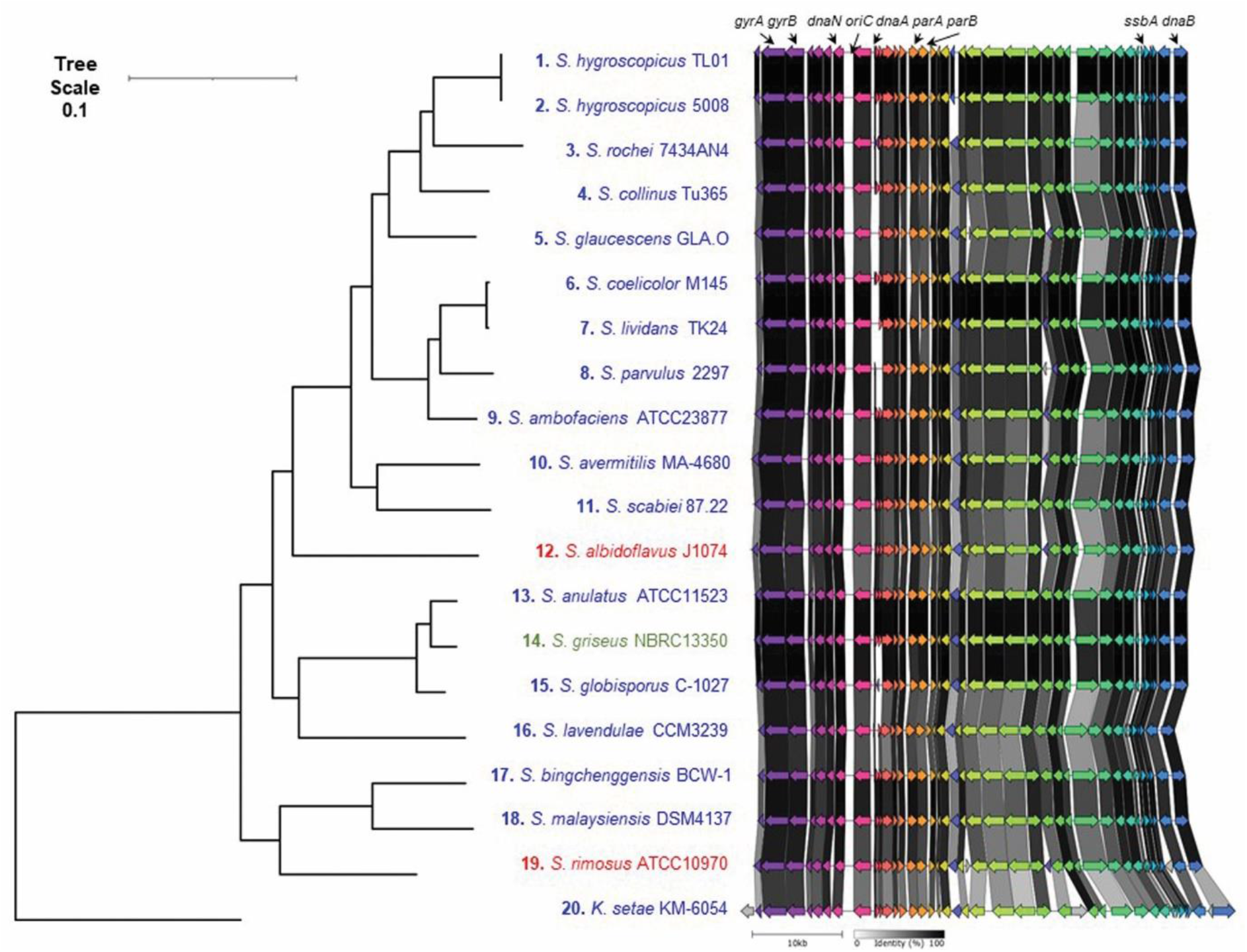
Streptomycete origin islands are syntenous. Origin islands from tRNA-ile (left) and *dnaB* (right*, SCO3911*) of 20 closed streptomycete genome sequences. Chromosomes with archetypal telomeres are listed in blue, Sg2247 telomeres in red and Sg13350 in green.

Bacterial chromosomes display a high degree of organization with respect to the locations of core genes (Lioy et al., 2021, Slager and Veening, 2016). In *Bordatella pertussis*, with a circular chromosome, there is a bias toward symmetrical inversions (Weigand et al., 2019) for the maintenance of replichore balance and preservation of favourable gene arrangements. Core-gene-defined genome organizational frameworks (cGOFs) can be classified as either symmetric or asymmetric with respect to their orientation in relation to the origin-terminus axis (Kang et al., 2014); in Gram-positive organisms cGOFs are exclusively symmetric and reversible in orientation (Kang et al., 2014). As a result it seems that the bilateral symmetry of streptomycete chromosomal rearrangements is consistent with Gram positive bacteria with circular chromosomes and that a bias exists towards maintaining an approximate equality in the lengths of both replichores.

### Asymmetric distribution of *parS* sites in streptomycete chromosomes

The delineation of the streptomycete origin island (Fig. 6) showed that this ~50 kb region not only contained *oriC* between *dnaN* and *dnaA*, but also a high number of *parS* sites located to the right of *oriC*. ParB binds to these sites and facilitates chromosome partitioning and connection with the polarisome (Jakimowicz et al., 2002, Kois-Ostrowska et al., 2016). As a result, we analysed the location of *parS* sites across the 20 closed streptomycete chromosomes. To do this, we employed a consensus matrix developed to locate prokaryotic *parS* sites (Livny et al., 2007).

The locations of these sites within the core region of the chromosome and with respect to *oriC* are shown in Fig. 7 and across the entire chromosome in Fig. S8. The location of each individual *parS* site is further described in Table S4. On average the number of *parS* sites per chromosome was 22.8 (range 11-65); most bacteria contain less than 5 *parS* sites (Livny et al., 2007), so the fact that streptomycetes carry up to 65 *parS* sites is remarkable (*S. albidoflavus* J1074 (Zaburannyi et al., 2014)). This strain displays fast and dispersed growth in liquid culture (Zaburannyi et al., 2014); it may be that the 65 predicted *parS* sites found in this strains contribute towards this through more efficient chromosome segregation and branching. As in *S. coelicolor* (Jakimowicz et al., 2002), most *parS* sites are located close to *oriC* in all strains and the origin island is abundant with *parS* sites; on average 45.24 % of *parS* sites are located within the origin island with a mean distance of 5,602 bp between *parS* sites in the origin island and 1,113,078 bp between *parS* sites in the bulk chromosome. Significantly, there is also a bias in the location of *parS* sites towards the right hand side of the chromosome (Fig. 7). 20.14% of *parS* sites lie to the left of *oriC* and 79.86% to the right (Table S4) meaning that during DNA replication and partitioning it is likely that the right replichore carries a greater abundance of ParB/*parS* nucleoprotein complexes with potential implications for chromosome segregation and connection of the chromosome to the hyphal tip. We did not include *Streptomyces venezuelae* 10712 in our analysis as the telomeres of its chromosome have not yet been established; although, the possession of Tap and Tpg suggests that they resemble the archetypal telomeres of *S. coelicolor* M145. However, ChIP-seq analysis confirmed the formation of large nucleoprotein ParB complexes located at 16 *parS* sites in *S. venezuelae* 10712 (Donczew et al., 2016) and recently, the role of ParB in promoting chromosome interarm contact was reported (Szafran et al., 2020). *parS* sites are also located to the right of *oriC* in the *S. venezuelae* chromosome (data not shown). The significance of this bias of *parS* sites on the right replichore is unclear; it is true that only one daughter chromosome is directed towards the hyphal tip by ParB in *S. coelicolor* (Kois-Ostrowska et al., 2016), but it is difficult to imagine how a bias in ParB binding to the right replichore would manifest itself in the preferential direction of one daughter chromosome toward the hyphal tip.

**Fig. 7.**
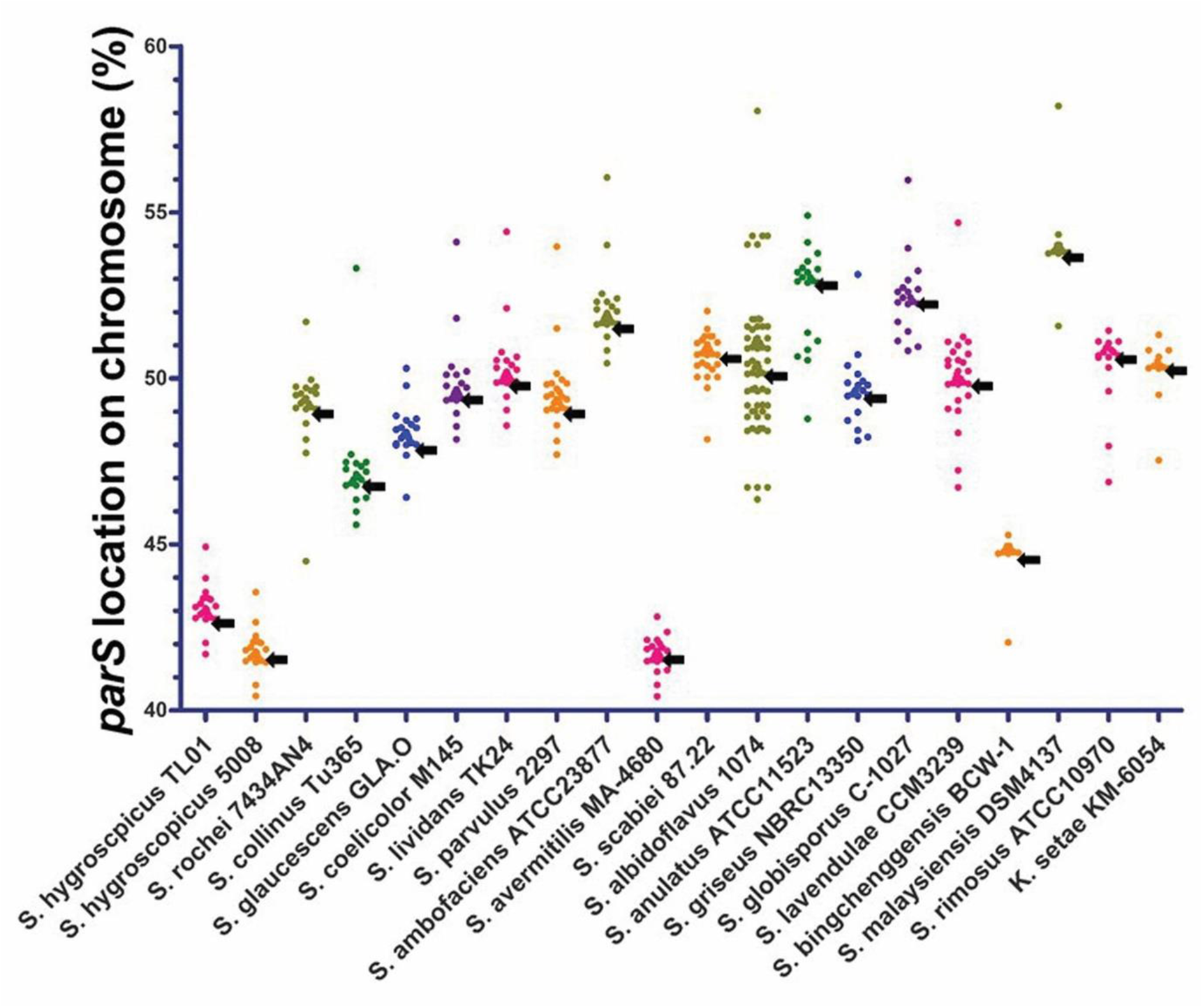
Location of *parS* sites in the central 20% of streptomycete chromosomes. The location of predicted *parS* sites on the chromosomes of the 20 closed streptomycete genomes were determined by interrogation of those genomes with the consensus matrix for bacterial *parS* sites (Livny et al., 2007). Locations were expressed as a percentage of the chromosome size. The last base of *dnaA* from all genomes was selected as the location of *oriC* (black arrows).

In summary, following the completion of the *S. rimosus* ATCC 10790 genome sequence, the parental strain of an important lineage of industrial bacteria, we first characterised an uncommon set of telomeres for this organism before comparing them to the other five classes of streptomycete replicon ends. Through the organisation of the closed replicons of 20 streptomycete chromosomes with respect to their phylogeny and physical orientation we determined that the telomeres were not associated with particular clades and were likely shared amongst different strains by horizontal gene transfer. We identified an origin island that forms an axis around which symmetrical chromosome inversions can take place and that there is a bias in *parS* sites to the right of *oriC* in closed streptomycete genomes. These results open up new questions for further investigations towards understanding the mechanisms of end patching of streptomycete chromosomes and the biological meaning that lies in the *parS* site bias. As such, understanding the fundamental structure of genome organisation and mechanisms will prove indispensable in strain engineering for improved specialised metabolite production by this industrially important microbial group.

### Methods

#### Genomic DNA extraction, Sequencing and Genome Assembly

*S. rimosus* was obtained from Pfizer (Petkovic et al., 2006) and propagated on Emerson’s Agar at 30°C (g/L: 4 Beef extract, 1 Yeast extract, 4 Peptone, 10 Dextrose, 2.5 NaCl, 20 Agar, pH 7.2) before inoculating TSB medium (Kieser et al., 2000), incubated at 30°C, 250 rpm in 250 ml baffled flasks for 36 hours prior to genomic DNA isolation or preparation of PFGE plugs. For the former, biomass was harvested by centrifugation and resuspended in 300 μl TE buffer, pH 7.5 with 10 mg/mL lysozyme and 0.1 mg/mL RNase and incubated for 90 min at 37°C. 50 μl of 10% (w/v) SDS was added and the sample mixed followed by adding 85 μl of 5M NaCl. 400 μl of phenol/chloroform/isoamyl alcohol (25:24:1) were added, vortexed for 30s and centrifuged for 10 min at 8000 rpm. The aqueous layer was added to a new tube and the previous two steps were repeated. 400 μl of chloroform/isoamyl alcohol (24:1) were added and vortexed for 30s and centrifuged for 10 min at 8000 rpm. The aqueous layer was added to a new tube and 0.5mL of isopropanol added. The tube was inverted and incubated 5 for min at room temperature. DNA was pelleted at 10000 rpm, washed with 70% (v/v) ethanol, the supernate removed and the pellet air-dried for 30 min at room temperature. The DNA was resuspended in 10 mM Tris buffer pH 7.5 and sent for sequencing by PacBio at Nu-omics (https://www.northumbria.ac.uk/business-services/engage-with-us/research/nu-omics/ and by Illumina 2 x 250bp paired-end reads at MicrobesNG (https://microbesng.com/). DNA was quantified using the Qubit dsDNA HS assay. Details of sequence assembly are provided in Supplementary information.

#### Determination of telomere sequences

Purification of the telomeric sequences was performed following an adaptation of a previously-described procedure (Fan et al., 2012). 1 μg of genomic DNA of *S. rimosus* was digested overnight with the blunt-end restriction enzymes SmaI and PvuII. The products were then purified using the Promega PCR-Clean Up kit and eluted in a final volume of 90 μl with dH_2_O. 10 μl of 1M NaOH was added and the mixture was incubated for 1 hour at 37°C. The samples were neutralised using 2M HCl and 1M Tris (pH 8) was added to a final concentration of 0.1M. 20X SSC solution was then added to final concentration of 2X and the samples were incubated at 68°C for 1 hour. The samples were purified once again with the Promega PCR-Clean Up kit and ligated overnight using Promega T_4_ DNA ligase. Inverted PCR analyses were performed using the primers listed in Table S1 and the amplified products were purified and sequenced by Eurofins Genomics.

#### Pulsed Field Gel Electrophoresis

PFGE analyses were performed using the Bio-Rad CHEF-DR^®^ II PFGE. Agarose plugs containing DNA were prepared from TSB grown liquid cultures of *S. rimosus* using established procedures (Kieser et al., 2000) with the following amendments to reduce DNA degradation (Evans and Dyson, 1993). HEPES was substituted for TRIS in buffers and mycelium was washed in HES buffer (25mM HEPES-NaOH, 25mM EDTA, 0.3M Sucrose pH8) and digested in 1 mg/ml lysozyme in HES buffer. After lysis, plugs were washed three times with NDS (1% N-laurylsarcosine, 0.5M EDTA, 10mM Glycine, pH9.5). The plugs were incubated overnight with NDS containing 1mg/ml Proteinase K and 1mM CaCl_2_. The next day, the plugs were washed in HE buffer (10mM HEPES-NaOH, 1mM EDTA, pH:8). After 15 minutes at room temperature, the liquid was replaced with fresh buffer and 1 μl of bovine serum albumin and 50 U of AseI or DraI added. The mix was incubated overnight at 37 °C. In case of AseI digestion, a second sample of 10 U of enzyme was added after 2 hours. For electrophoresis, HEPES buffer (16mM HEPES-NaOH, 16mM Sodium acetate, 0.8mM EDTA, pH7.5) was used and the voltage set at 4V/cm, with an initial switch time of 70 seconds and a final switch time of 130 seconds. Eelectrophoresis was performed for 24 hours. The gel was then stained for 30 minutes in 1 μg/ml ethidium bromide solution and de-stained with dH_2_O for an hour. The gel was imaged with a UV trans-illuminator.

#### Telomere identification and analysis

In order to identify streptomycete telomeres, we used the terminal 36 and 150bp of the six classes of telomeres to search NCBI using BLASTn. The two sizes of query sequences were taken from the archetypal end from the *S. coelicolor* chromosome (Sco) (Bentley et al., 2002); the non-archetypal end of *S. coelicolor* plasmid SCP1 (SCP1) (Huang et al., 2007); the chromosome end of *S. griseus* 13350 chromosome Sg13350 (Ohnishi et al., 2008); the chromosome end of *S. rimosus* ATCC10970 chromosome (Sg2247) (this work); the end of *Streptomyces* sp. 44030 plasmid pRL1(pRL1) (Zhang et al., 2006); the end of *Streptomyces* sp. 440414 plasmid pRL2 (pRL2) (Zhang et al., 2006). The query sequences are described in Fig. 1. Only those significant hits from complete whole genome sequencing projects that were either located at the end of a replicon or where sequences were physically recovered and sequenced were designated as telomeres.

A ClustalW alignment of 13 Sg2247 independent telomeres was used to map telomere evolutionary history and was inferred using the Maximum Likelihood method and the Tamura 3-parameter model (Tamura et al., 2006). The tree with the highest log likelihood (−1238.57) was used. Initial tree(s) for the heuristic search were obtained automatically by applying Neighbor-Join and BioNJ algorithms to a matrix of pairwise distances estimated using the Maximum Composite Likelihood (MCL) approach, and then selecting the topology with superior log likelihood value. A discrete Gamma distribution was used to model evolutionary rate differences among sites (5 categories (+G, parameter = 0.6336)). Trees were drawn to scale, with branch lengths measured by the number of substitutions per site. This analysis involved 13 nucleotide sequences and there were a total of 163 positions in the final dataset. Evolutionary analyses were conducted in MEGA X (Kumar et al., 2018)

Stem-loop structures, identified using Mfold (Zuker, 2003), are displayed where the revised free energies (ΔG, kJ/mol) were determined using Jacobson-Stockmeyer theory to assign free energies to multi-branch loops. Projections were calculated using default conditions except folding temperature was set at 30°C, Na^+^ concentration of 0.05M and maximum distance between paired bases was set at 20 (Yang et al., 2017). Closed chromosome sequences from the genus *Streptomyces*, in conjunction with the closed sequence of *Kitasatospora setae* KM-6054 as an outgroup, were used to carry out Multi-Locus Sequence Analysis (MLSA) to produce a high-resolution species tree using AutoMLST after a concatenated alignment (Alanjary et al., 2019). The average nucleotide identities (ANI) between 20 closed streptomycete genomes was calculated using the OrthoANI tool (Lee et al., 2016) and plotted with the heat map function in R (Ver. 4.0).

Closed streptomycete chromosomes were subjected to a progressive alignment in Mauve (Darling et al., 2010) using default settings and dot-plots generated by Nucmer (Galaxy Version 4.0.0beta2+galaxy0) (Marcais et al., 2018) with default settings, except that the filter setting was turned on so only delta alignments which represent the ‘best’ hit to any particular location on either sequence were displayed. Maps showing synteny or origin islands were created using clinker & clustermap.js (Gilchrist and Chooi, 2020).

The location of predicted *parS* sites were determined by searching the 20 closed genomes using the consensus matrix for bacterial *parS* sites (Livny et al., 2007). This was done by performing a ClustalW alignment of the 1030 predicted *parS* sited previously identified (Livny et al., 2007). A consensus matrix was then constructed in Weblogo (http://weblogo.threeplusone.com/create.cgi) (Crooks et al., 2004) and used to interrogate closed streptomycete genomes using matrix scan in the Regulatory Sequence Analysis Tools suite (RSAT) (http://embnet.ccg.unam.mx/rsat//matrix-scan-quick_form.cgi) (Turatsinze et al., 2008). For this analysis the threshold weight score was set at >15 (Livny et al., 2007).

## Contributions

LAG and JKS isolated and purified *S. rimosus* genomic and telomere DNA. LAG, DRM and PRH carried out the bioinformatic analysis. ISH provided *S. rimosus* and contributed to intellectual input. RPH wrote the manuscript with the help of all co-authors. All authors discussed the results and commented on the manuscript. PRH supervised the study.

