## Supplementary Figs and Files for "Bilateral symmetry of linear streptomycete chromosomes": Bilateral Xsomes Supp Figs Final.pdf

<u>**Supplementary Figures and Tables**</u>

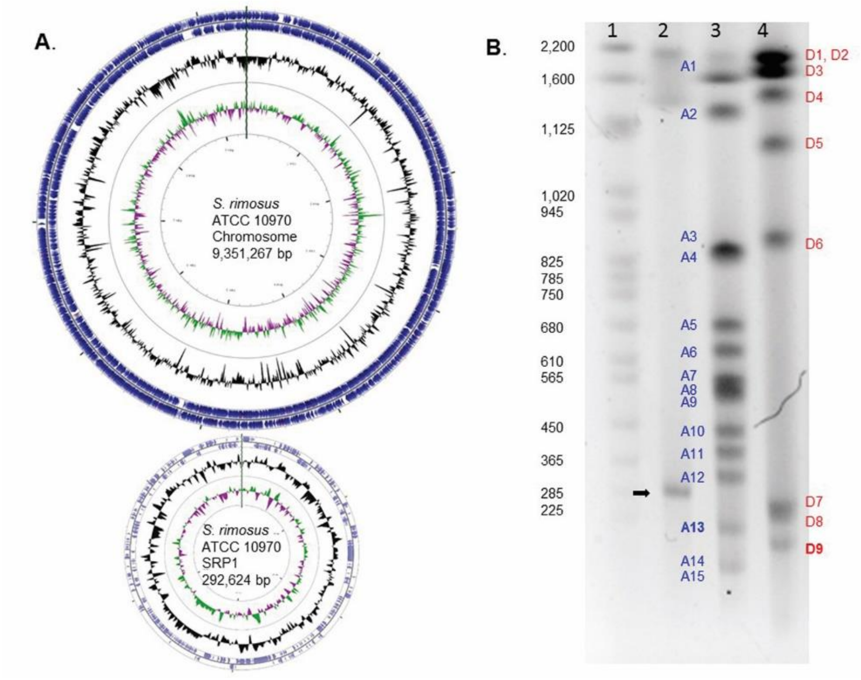


| AseI Fragment | Size (bp) | DraI Fragment | Size (bp) |
|---|---|---|---|
| A1 | 2,268,128 | D1 | 2,162,072 |
| A2 | 1,136,413 | D2 | 2,029,716 |
| A3 | 831,533 | D3 | 1,563,320 |
| A4 | 827,548 | D4 | 1,184,854 |
| A5 | 682,802 | D5 | 1,052,419 |
| A6 | 623,988 | D6 | 859,144 |
| A7 | 551,863 | D7 | 263,506 |
| A8 | 527,971 | D8 | 236,236 |
| A9 | 513,659 | **D9** | **172,852** |
| A10 | 433,169 | **D10** | **43,122** |
| A11 | 382,749 | **D11** | **30,090** |
| A12 | 320,709 | **D12** | **25,966** |
| **A13** | **207,231** | **D13** | **19,753** |
| A14 | 127,494 | **D14** | **841** |
| A15 | 123,241 | | |
| **A16** | **59,216** | | |
| **A17** | **26,177** | | |

**Fig. S1. The closed genome sequences of the linear chromosome and giant linear plasmid, SRP1, from *S. rimosus* ATCC10970. (A)** The linear replicons are represented as circles with the telomeres at 12 o'clock and *oriC* at 6 o'clock. From outside to inside, the concentric circles represent: nucleotide position; coding sequences (CDS) on the forward strand; CDS on the reverse strand; GC content (black); GC-skew on forward strand (purple); GC skew on the reverse strand (green). The genome sequence is listed in NCBI under accession number CP048261**. (B)** Pulsed Field Gel Electrophoresis analysis of *S. rimosus* ATCC10970. The size of SRP1 (lane 2, black arrow) was verified by Pulsed Field Gel Electrophoresis in comparison to yeast chromosomal markers (Biorad, lane 1). Restriction digestion with AseI (lane 3) and DraI (lane 4) of *S. rimosus* mycelial plugs followed by PFGE, allowed us to corroborate the genome assembly by comparison of the observed fragments with the *in silico* predictions of band sizes.

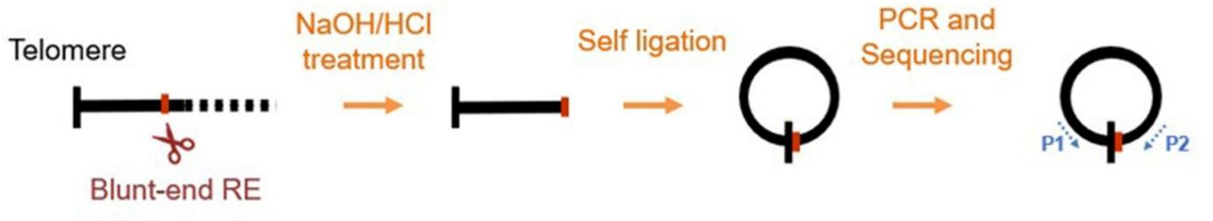


**Fig. S2. Scheme for recovery of *S. rimosus* telomeres.** In order to complete the genome sequence of *S. rimosus* we employed self-ligation of blunt ends coupled with inverted PCR (primers P1 and P2) and sequencing of the resulting amplicons (Fan et al., 2012) to recover the telomeres of the chromosome and SRP1.

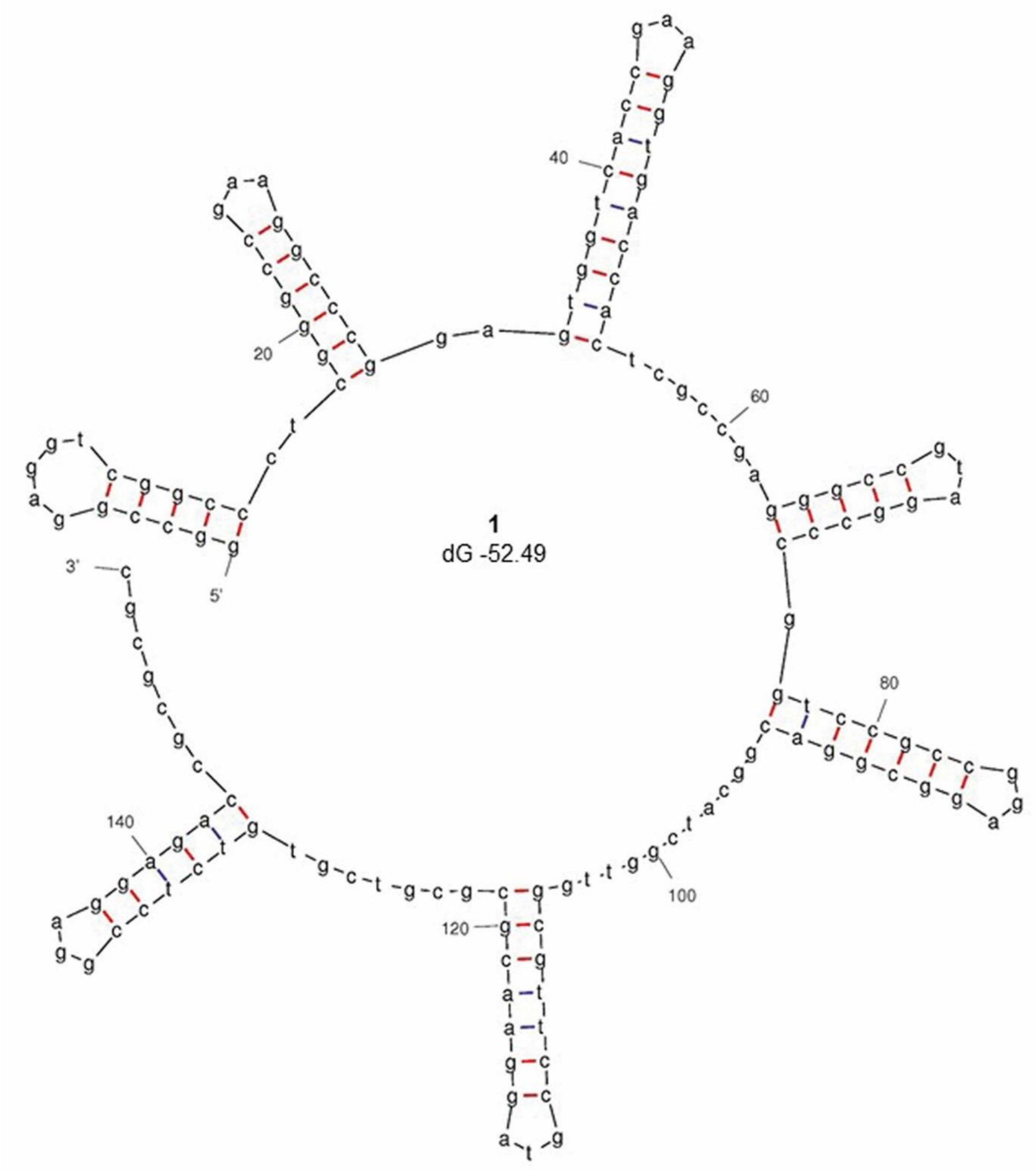


*S. albidofavus* J1074 chromosome, accession number NC_020990.1

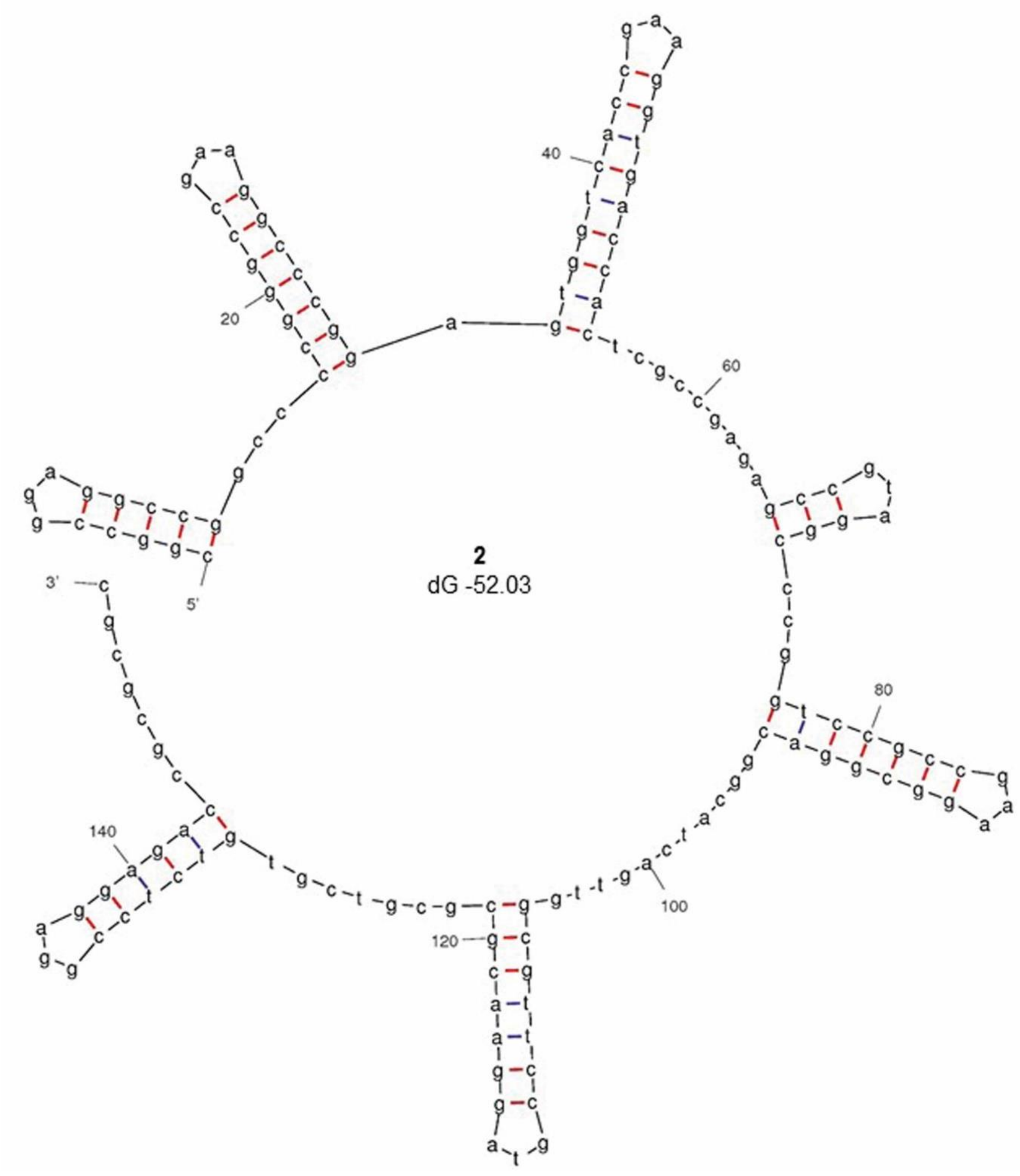


*Streptomyces* sp. HK1 plasmid pSHK1, EU372836.1

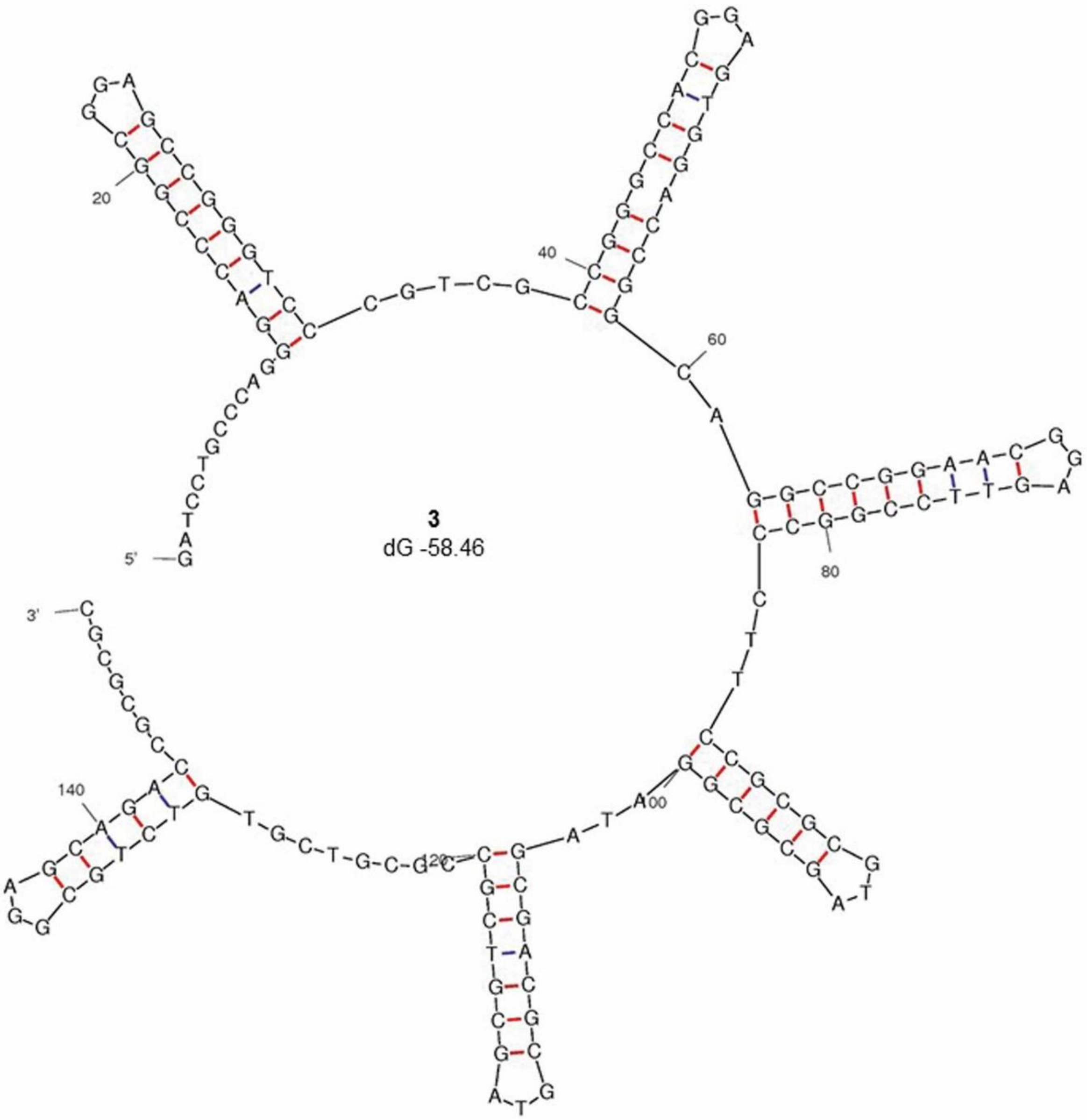


*Streptomyces* sp. 36R-2-1B plasmid pYY8L, GU080325.1

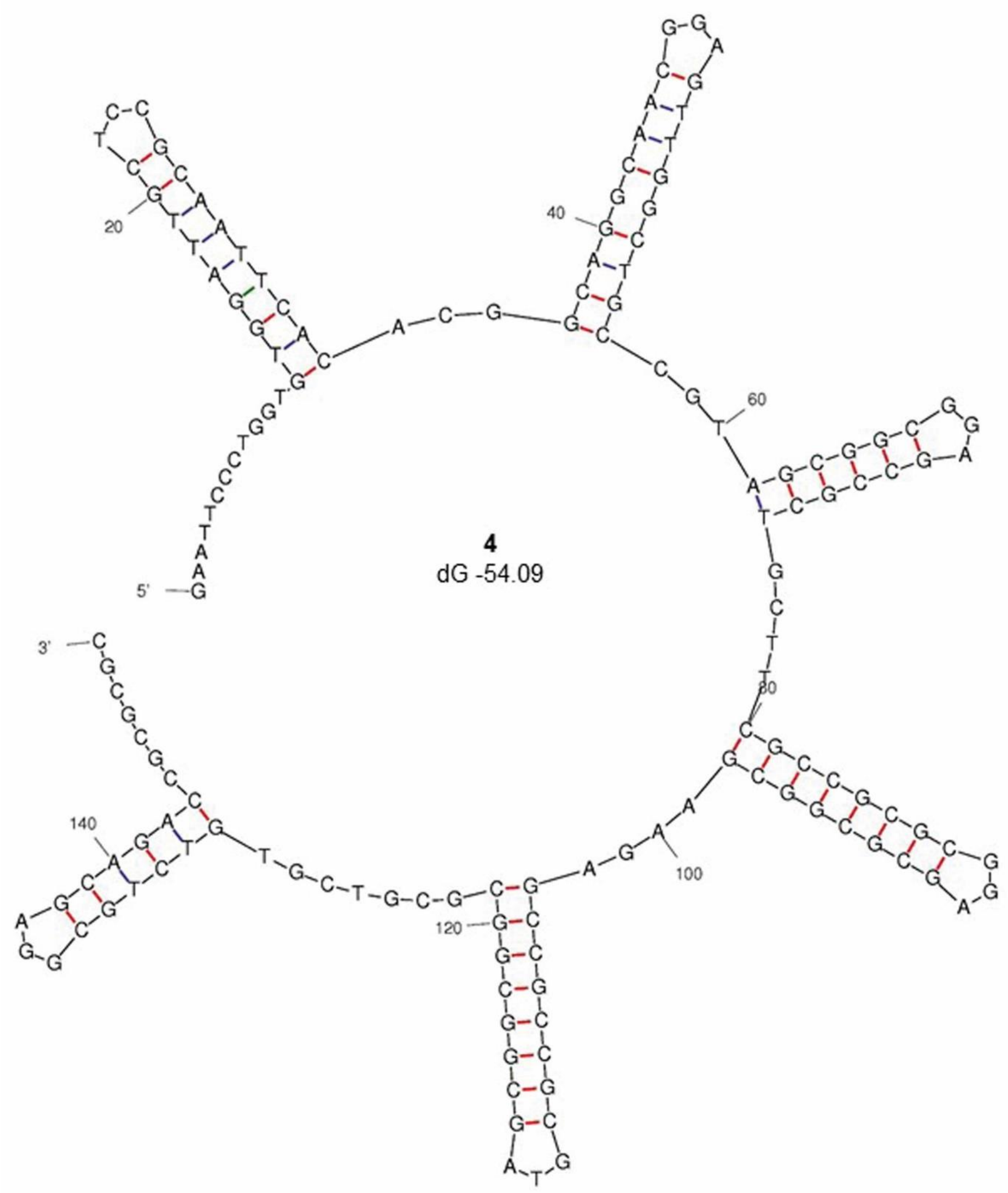


*S. cattleya* NRRL 8057 chromosome, FQ859185.1

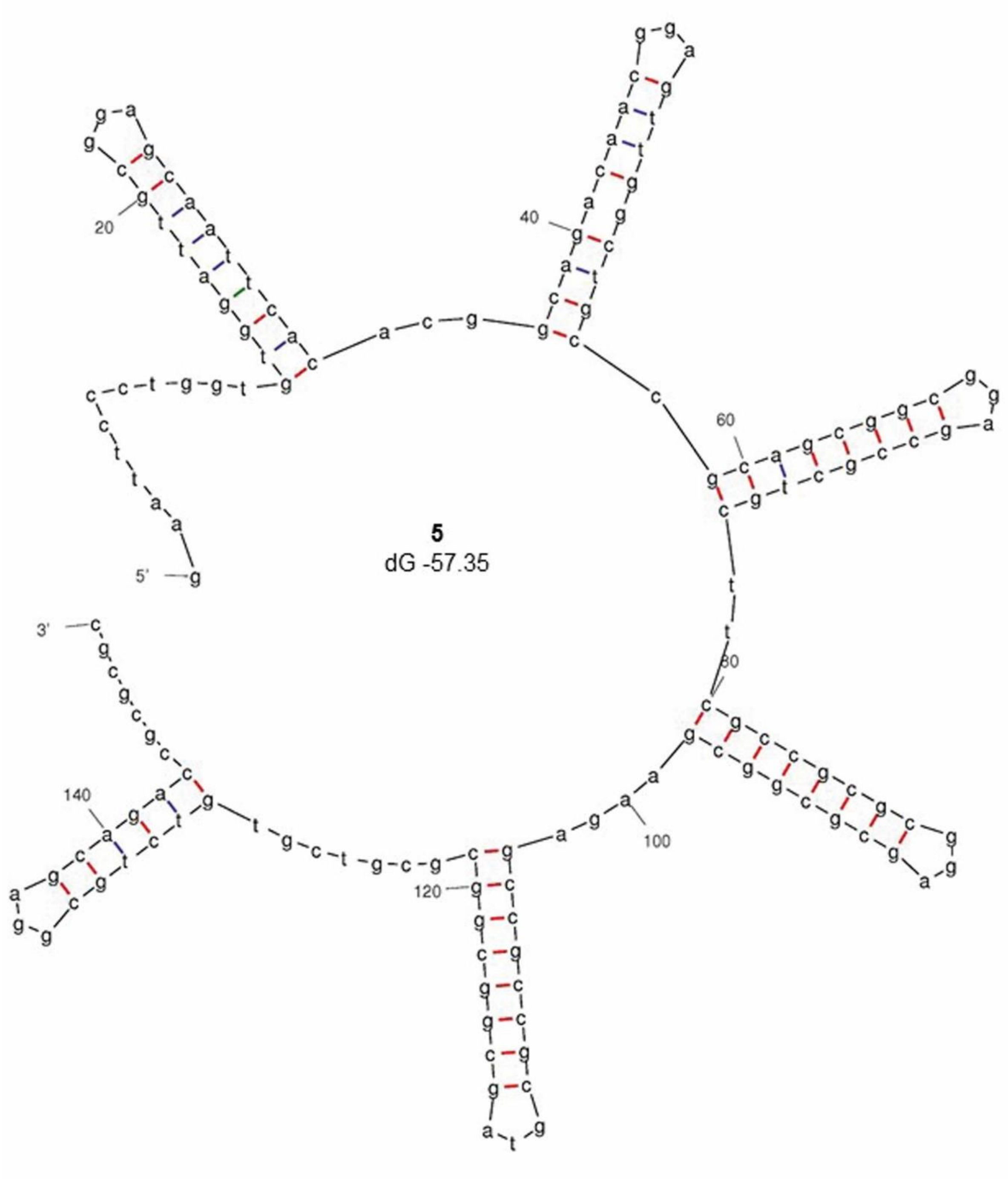


*S. cattleya* NRRL 8057 plasmid pSCAT, FQ859184.1

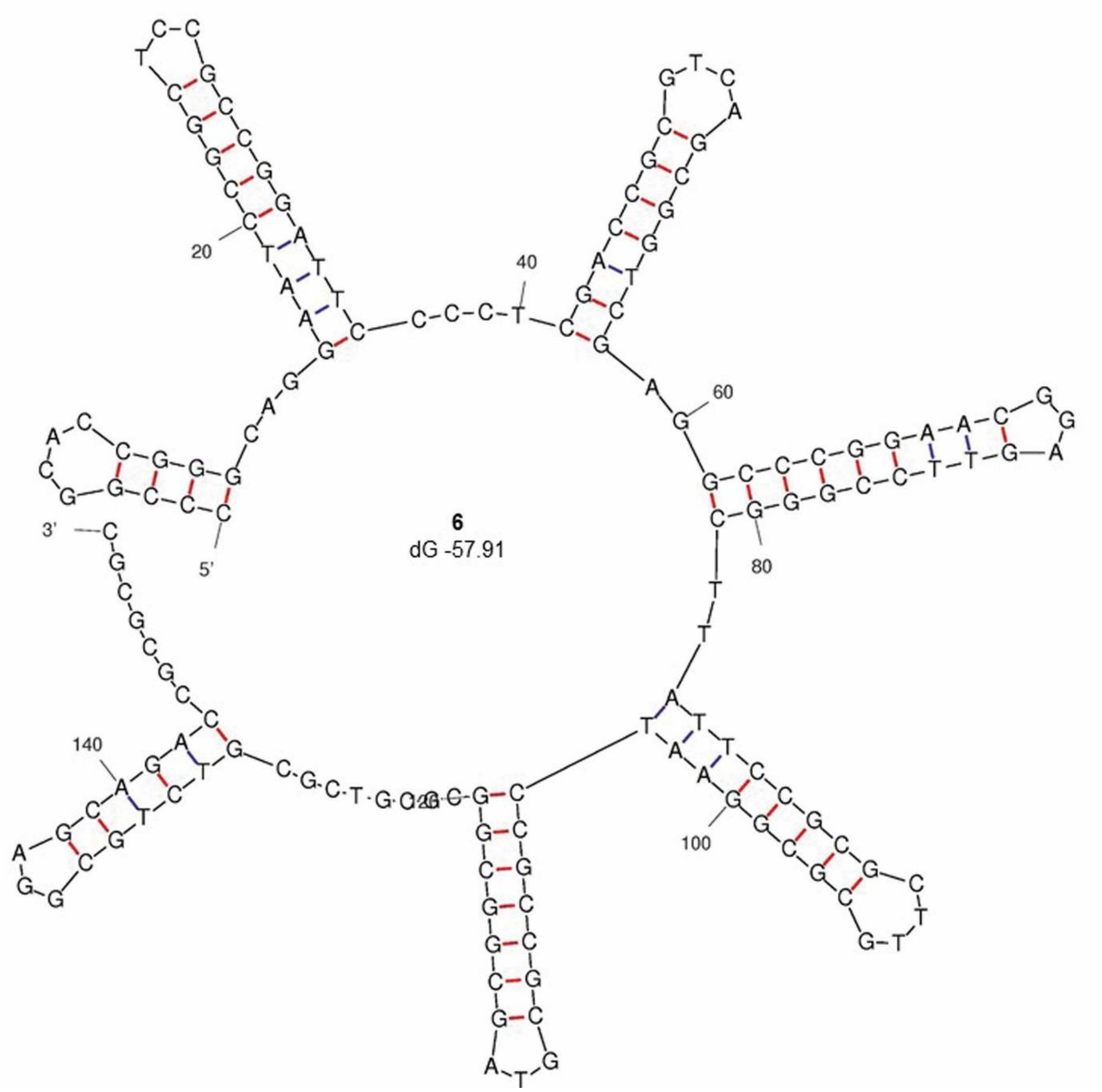


*S. hygroscopicus* subsp. jinggangensis TL01 plasmid pSHJGH1 right hand end, NC_020894.1

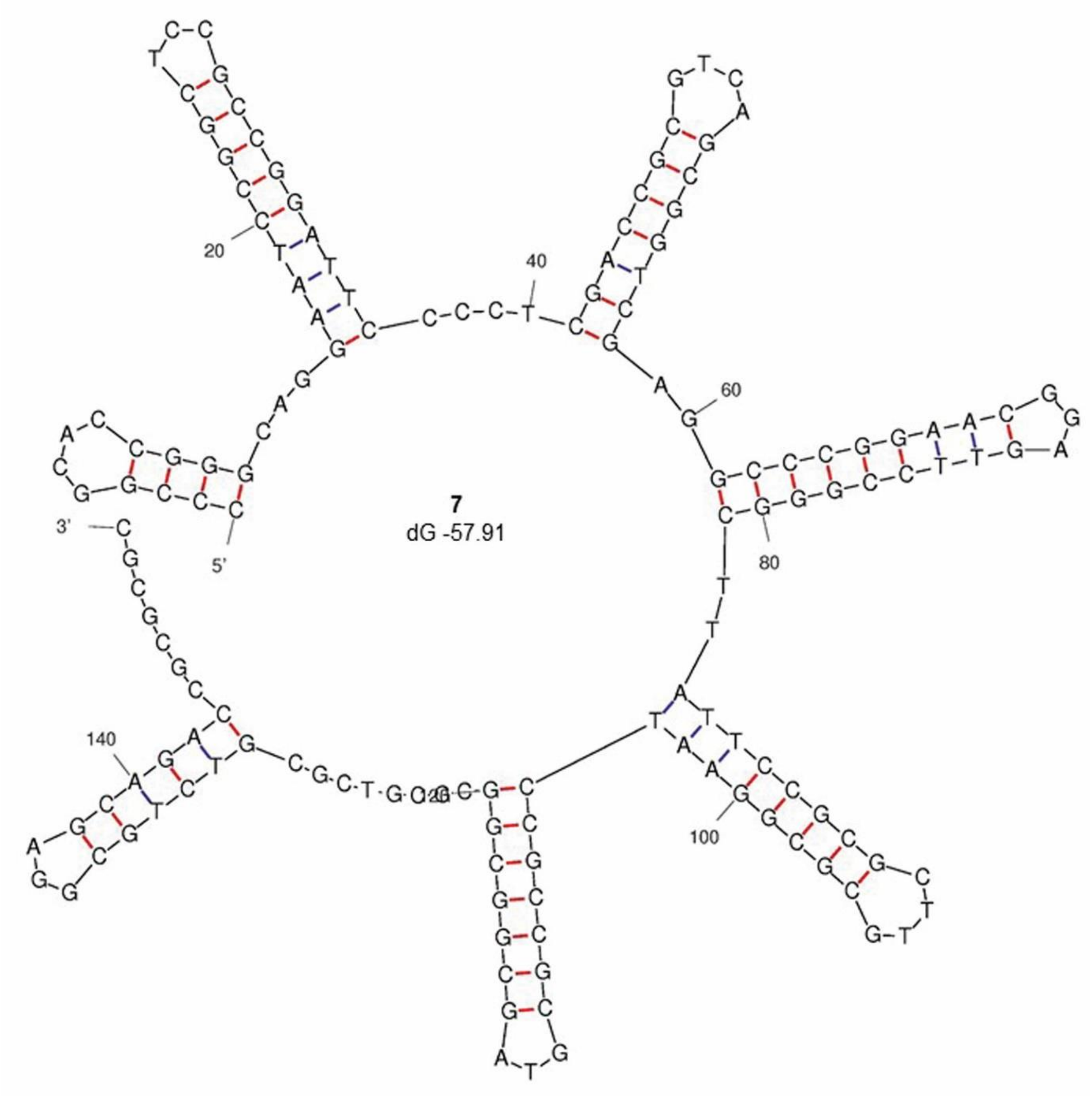


*S. hygroscopicus* subsp. jinggangensis 5008 plasmid pSHJG1 right hand end, NC_017766.1

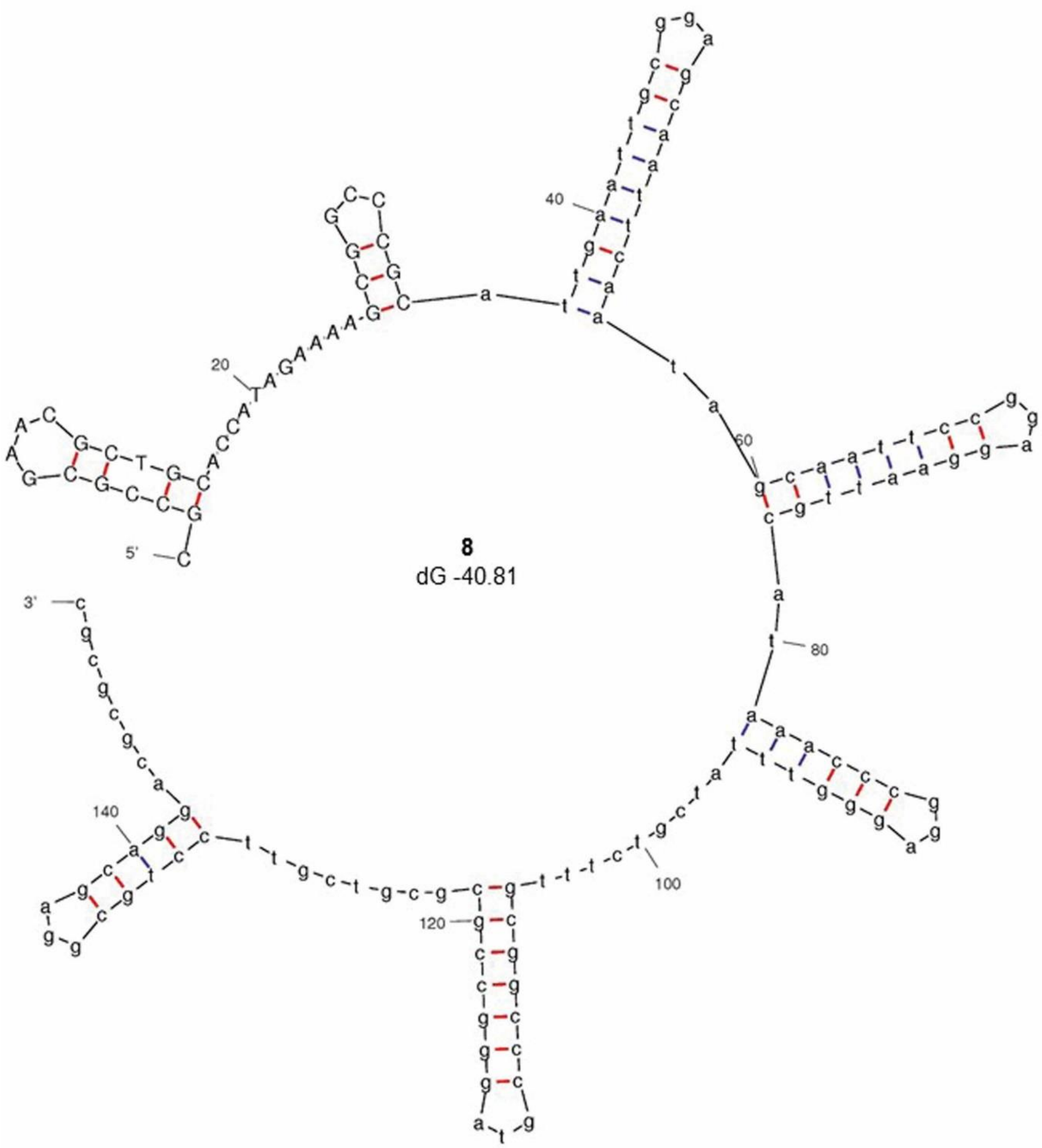


*S. rimosus* ATCC10970 chromosome, CP048261.1

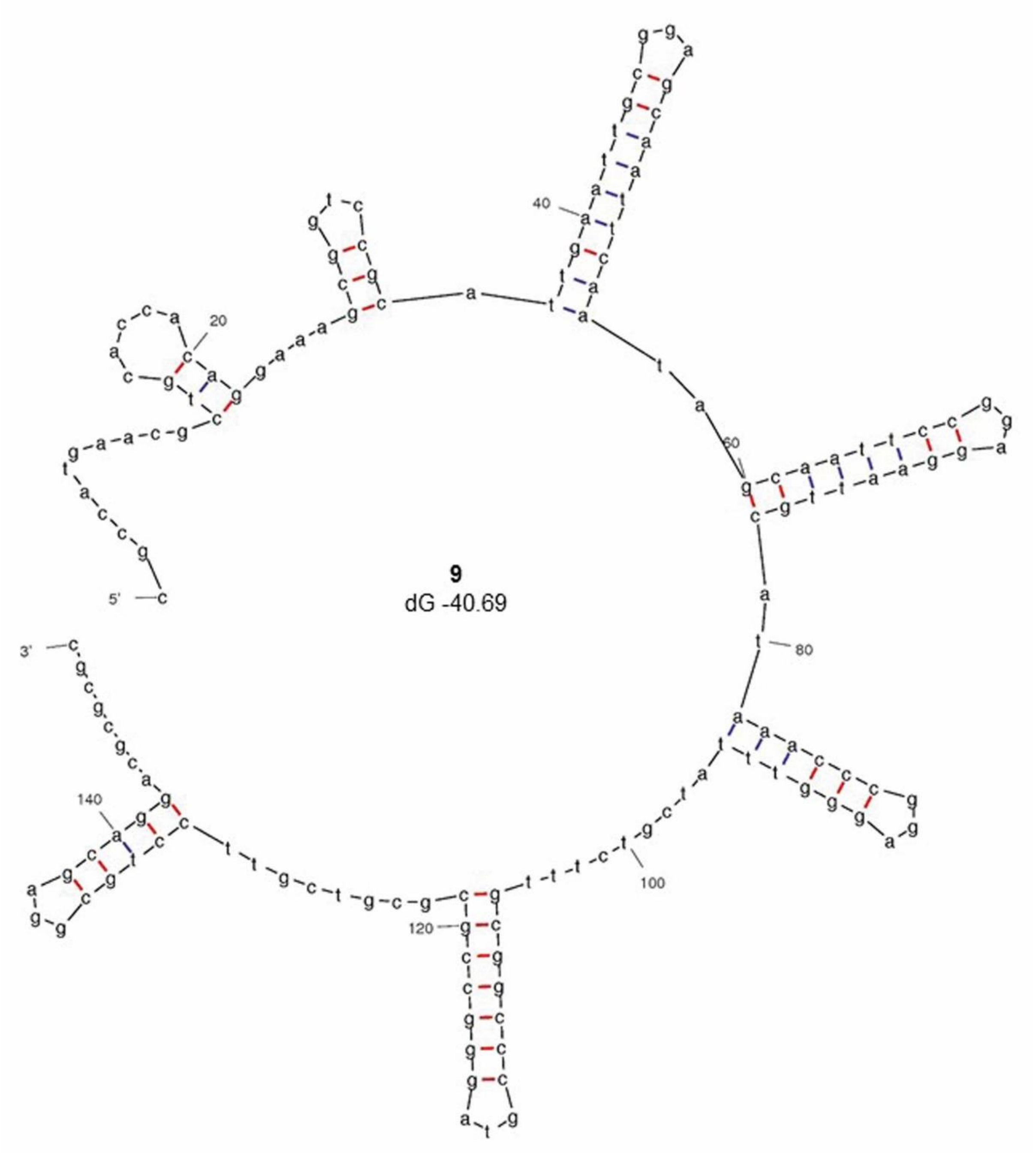


*S. rimosus* ATCC10970 plasmid SRP1 left hand end, CP048261.2

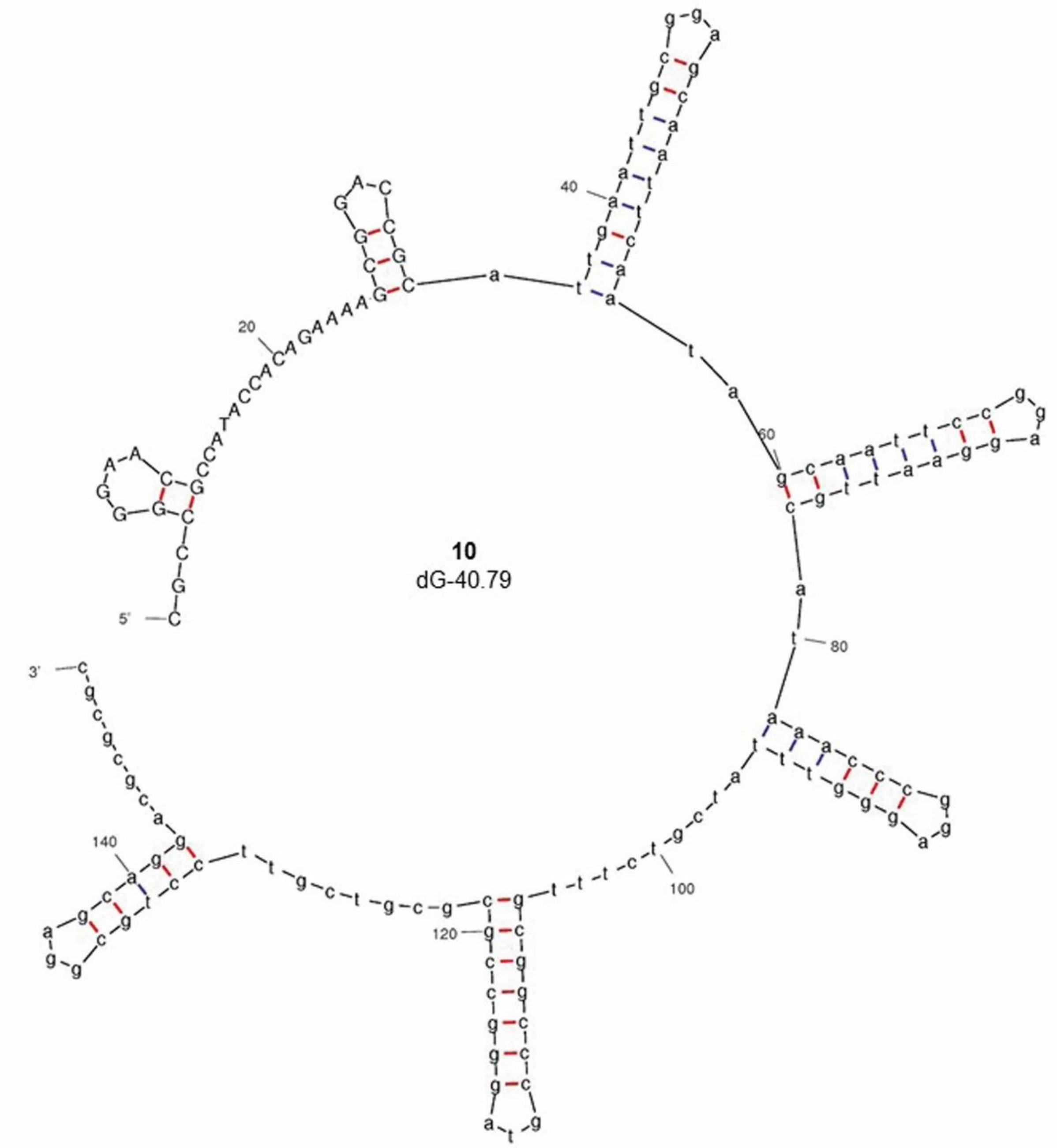


*S. rimosus* ATCC10970 plasmid SRP1 right hand end, CP048261.2

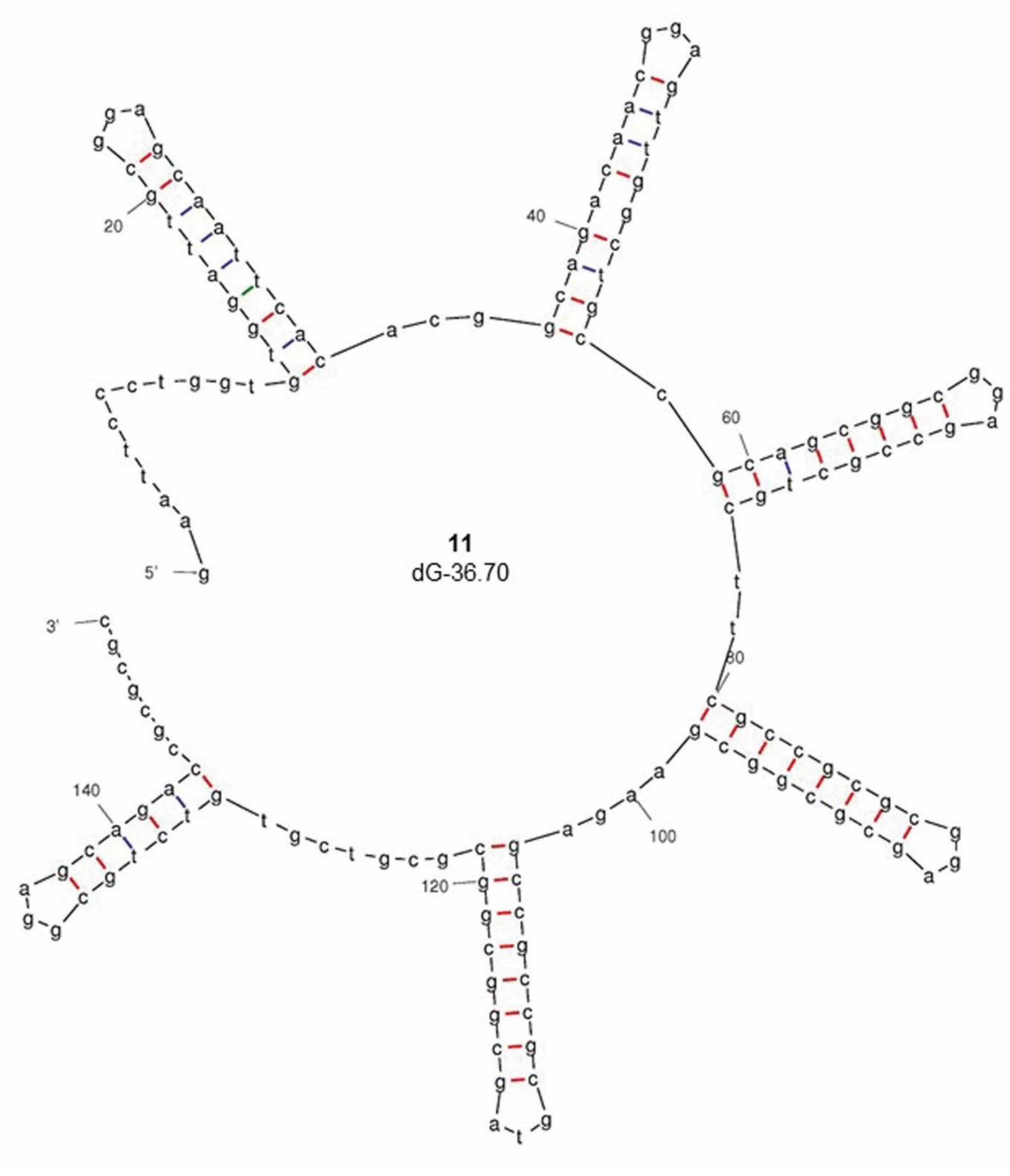


*S. pratensis* ATCC 33331 plasmid pSFLAO1, CP002476.1

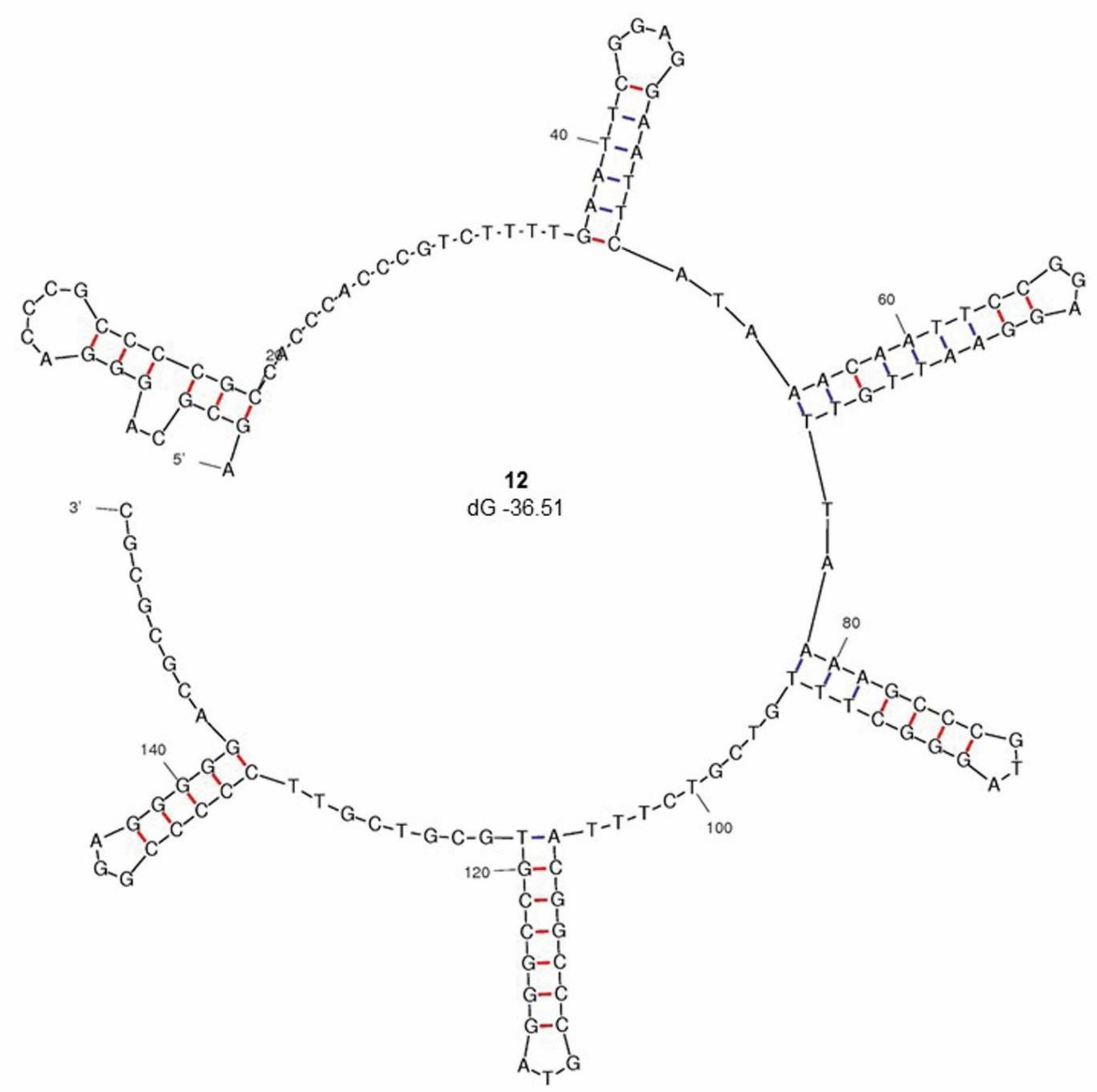


*S. griseus* 2247 chromosome

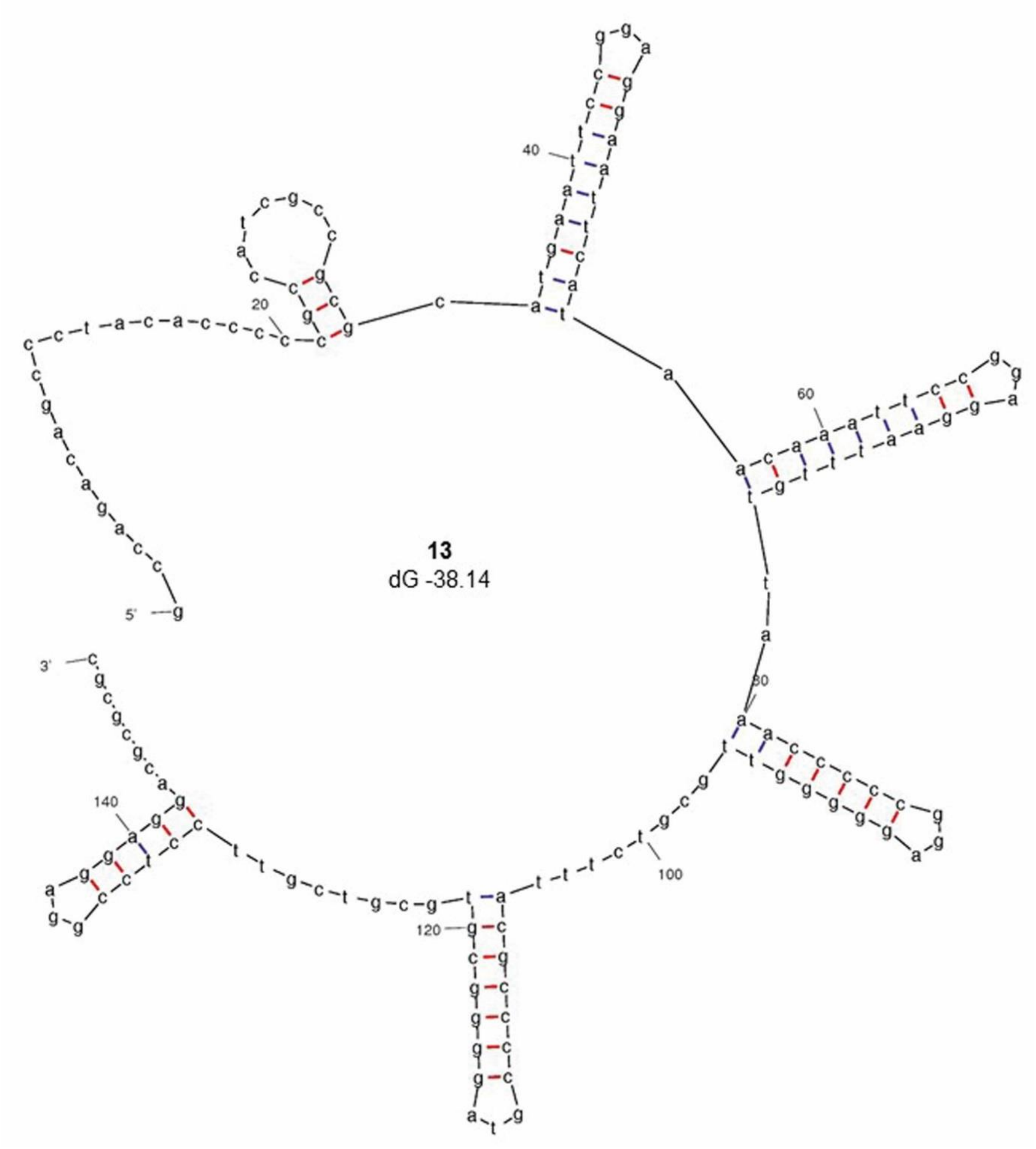


*Streptomyces* sp. 769 plasmid pSGZL, CP003988.1

**Fig. S3. Mfold projections of Sg2247-type telomeres.** The terminal 150bp of 3'replicon ends of Sg2247-type telomeres showing stem-loop structures and hairpins (I-VIII) identified using Mfold (Zuker, 2003) are displayed where the revised free energies (ΔG, kJ/mol) were determined using Jacobson-Stockmeyer theory to assign free energies to multi-branch loops. Projections were calculated using default conditions except folding temperature was set at

30°C, $Na^+$ concentration of 0.05M and maximum distance between paired bases was set at 20 (Yang et al., 2017). Further details of the 13 Sg2247-type telomeres are described in Table S2. 1, *S. albidofavus* J1074 chromosome, accession number NC_020990.1; 2, *Streptomyces* sp. HK1 plasmid pSHK1, EU372836.1; 3, *Streptomyces* sp. 36R-2-1B plasmid pYY8L, GU080325.1; 4, *S. cattleya* NRRL 8057 chromosome, FQ859185.1; 5, *S. cattleya* NRRL 8057 plasmid pSCAT, FQ859184.1; 6, *S. hygroscopicus* subsp. jinggangensis TL01 plasmid pSHJGH1 right hand end, NC_020894.1; 7, *S. hygroscopicus* subsp. jinggangensis 5008 plasmid pSHJG1 right hand end, NC_017766.1; 8, *S. rimosus* ATCC10970 chromosome, CP048261.1; 9, *S. rimosus* ATCC10970 plasmid SRP1 left hand end, CP048261.2; 10, *S. rimosus* ATCC10970 plasmid SRP1 right hand end, CP048261.2; 11, *S. pratensis* ATCC 33331 plasmid pSFLAO1, CP002476.1; 12, *S. griseus* 2247 chromosome; 13, *Streptomyces* sp. 769 plasmid pSGZL, CP003988.1. Further details of known streptomycete telomeres are listed in Table S2.

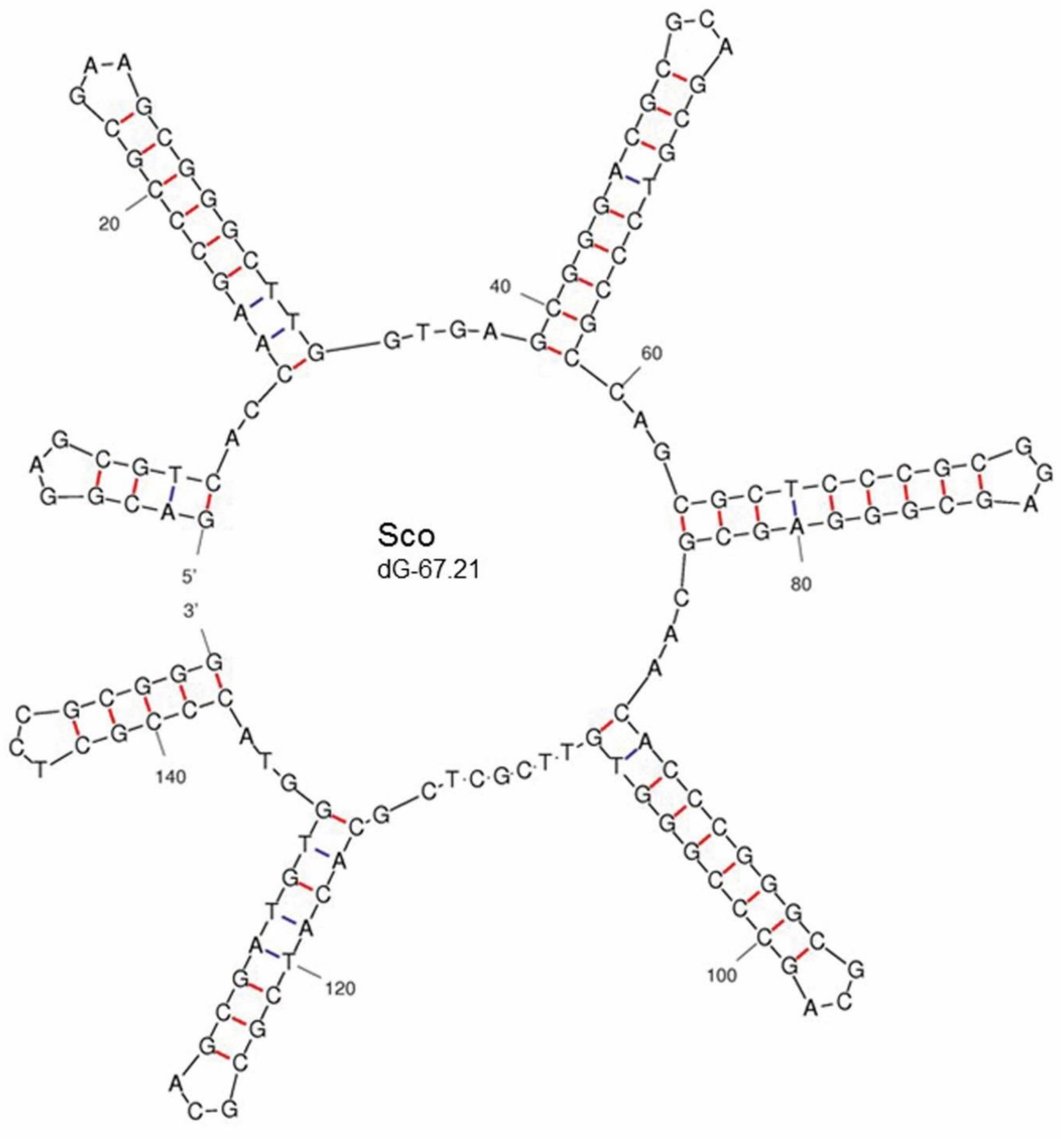
Sco
dG-67.21
5'
3'
20
40
60
80
100
120
140

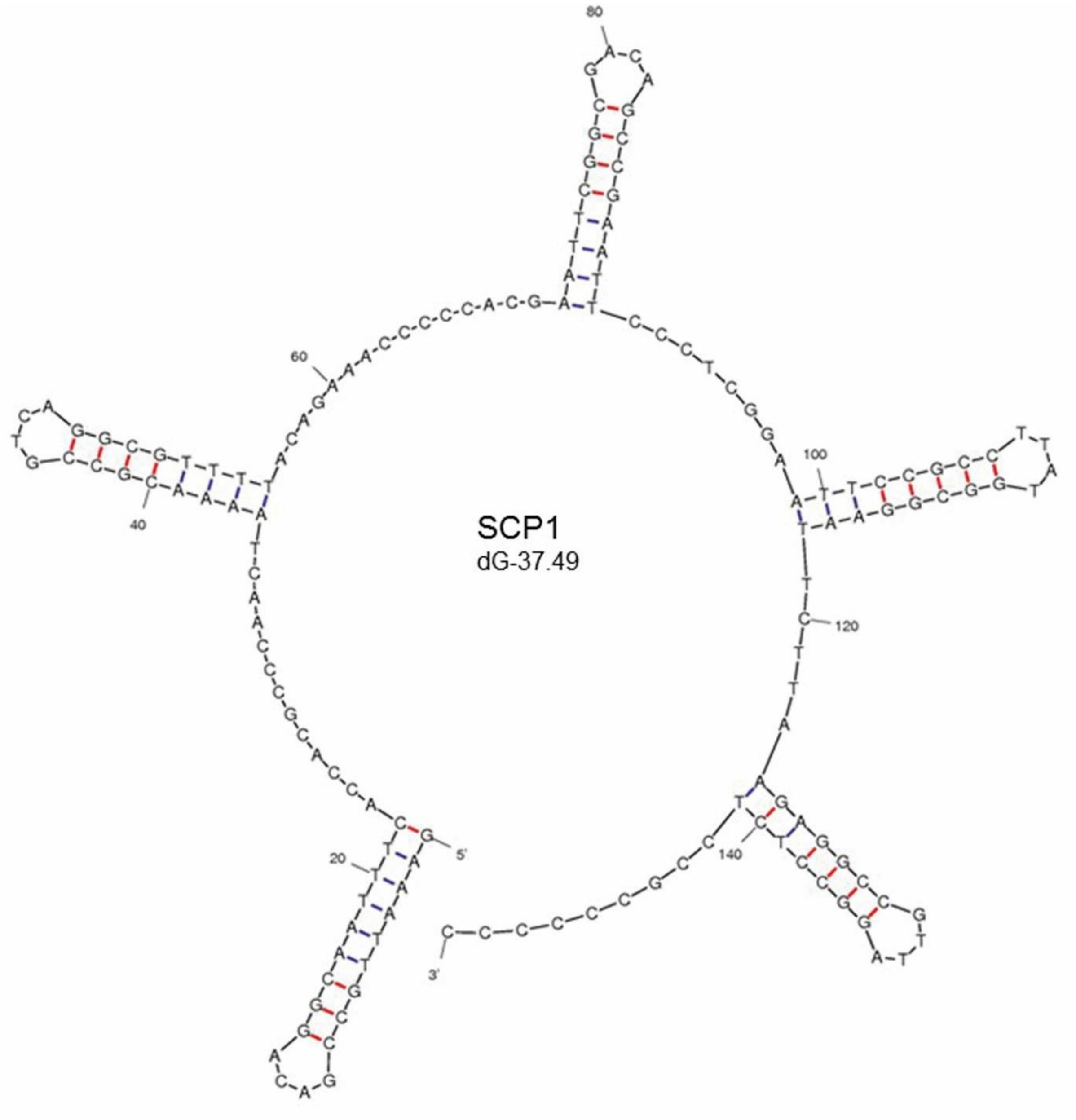
SCP1
dG-37.49
80
60
100
40
120
20
140
5'
3'

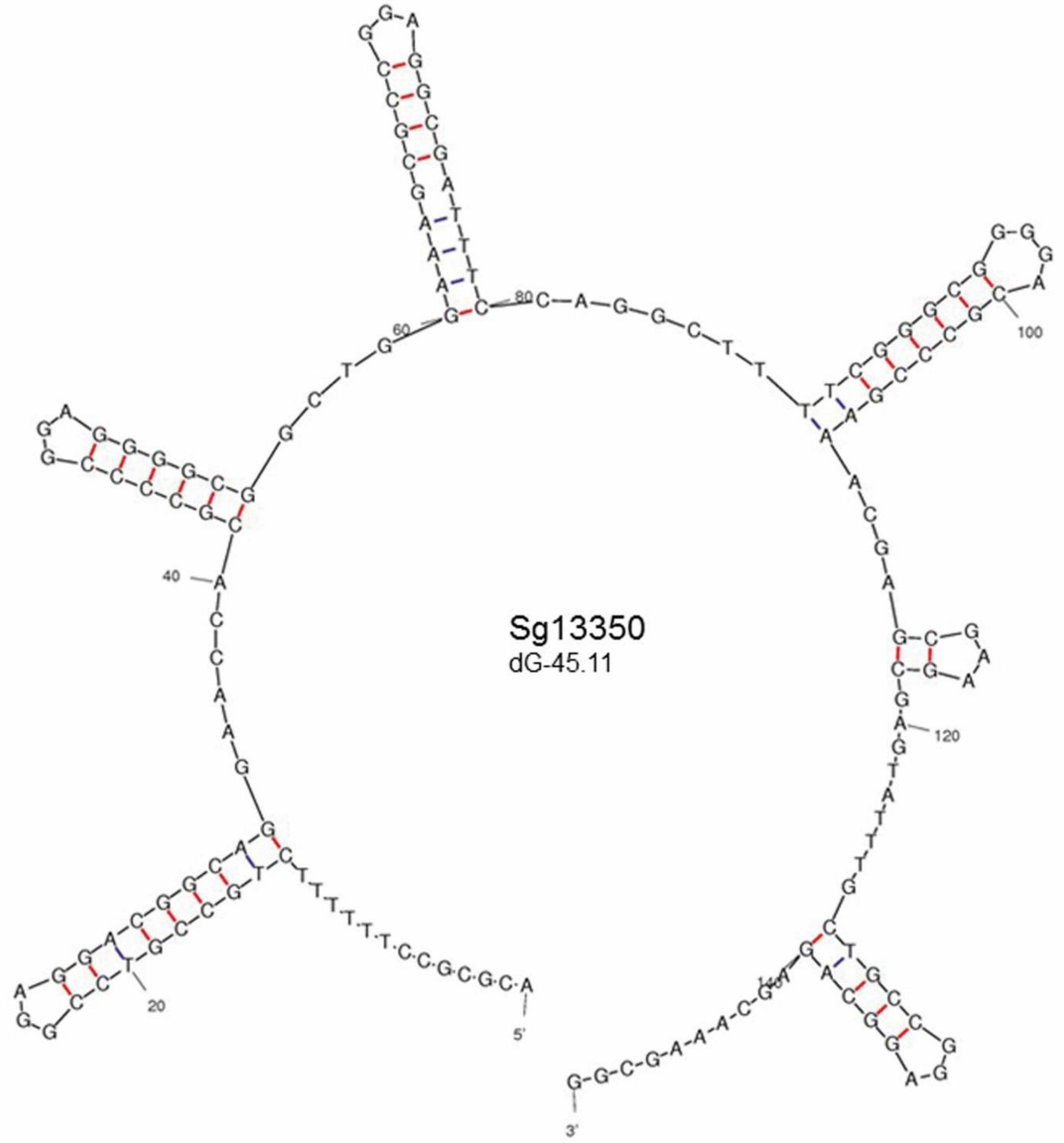
Sg13350
dG-45.11
5'
3'
20
40
60
80
100
120

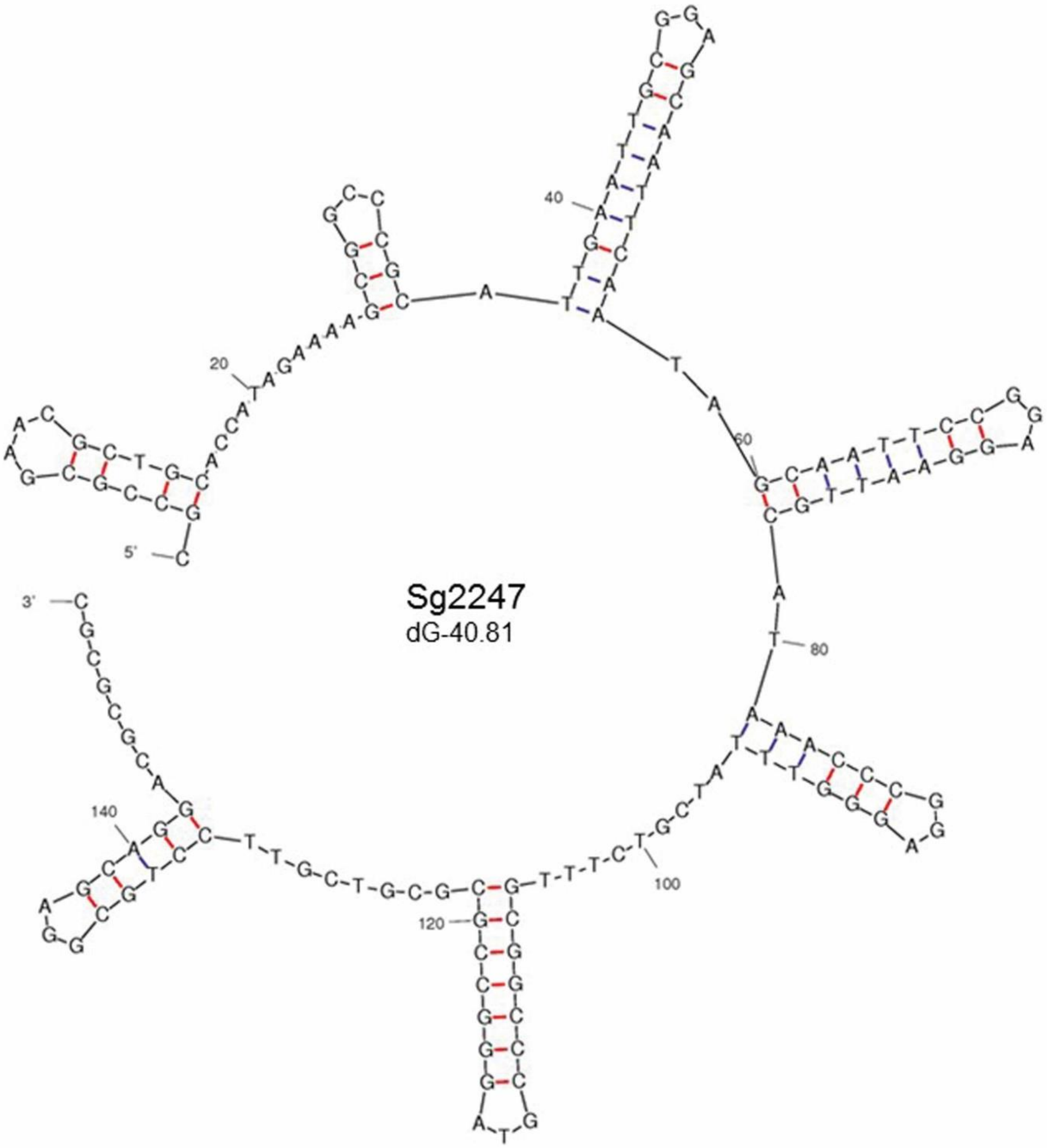
Sg2247
dG-40.81
5'
3'
20
40
60
80
100
120
140

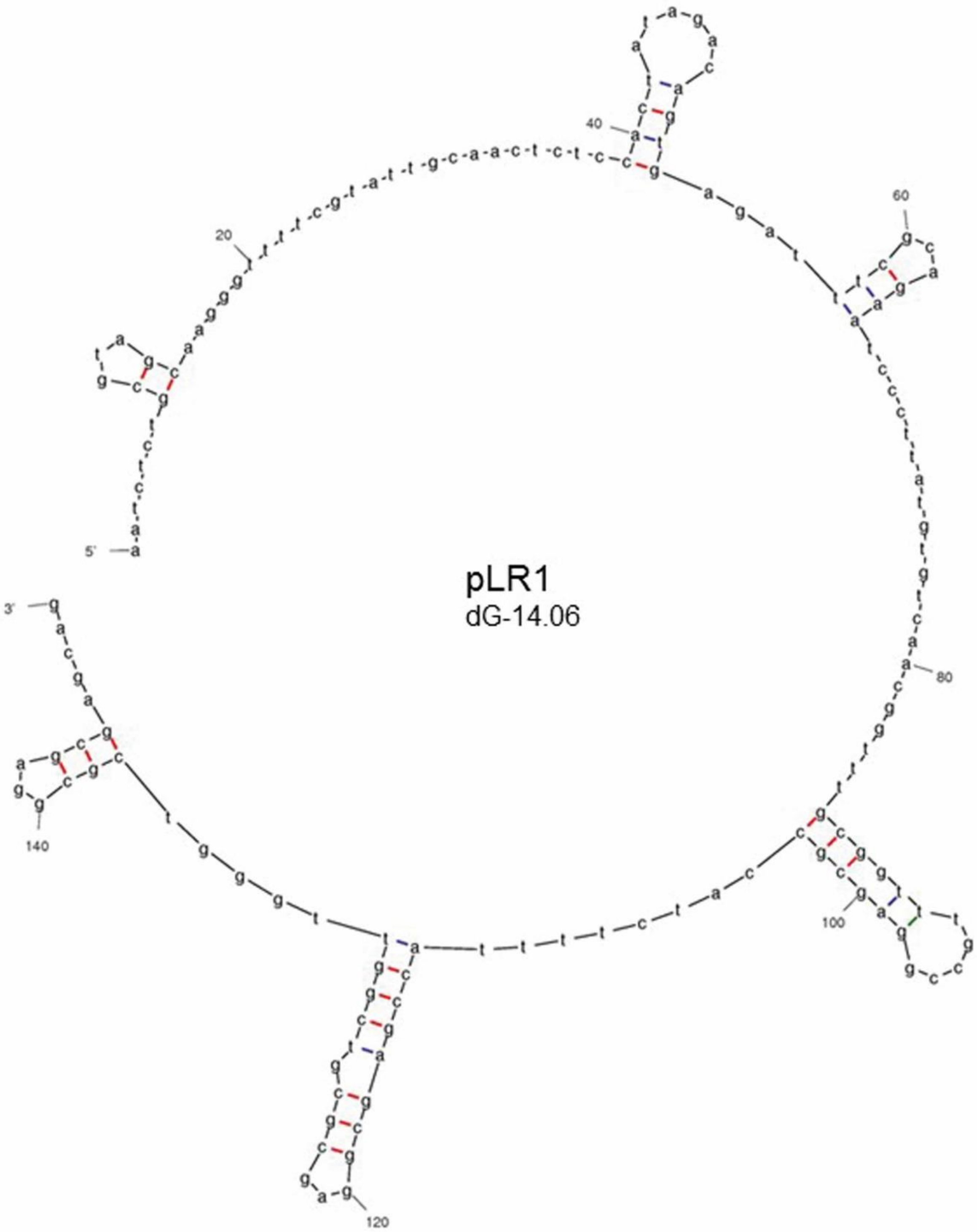
pLR1
dG-14.06
5'
3'
20
40
60
80
100
120
140

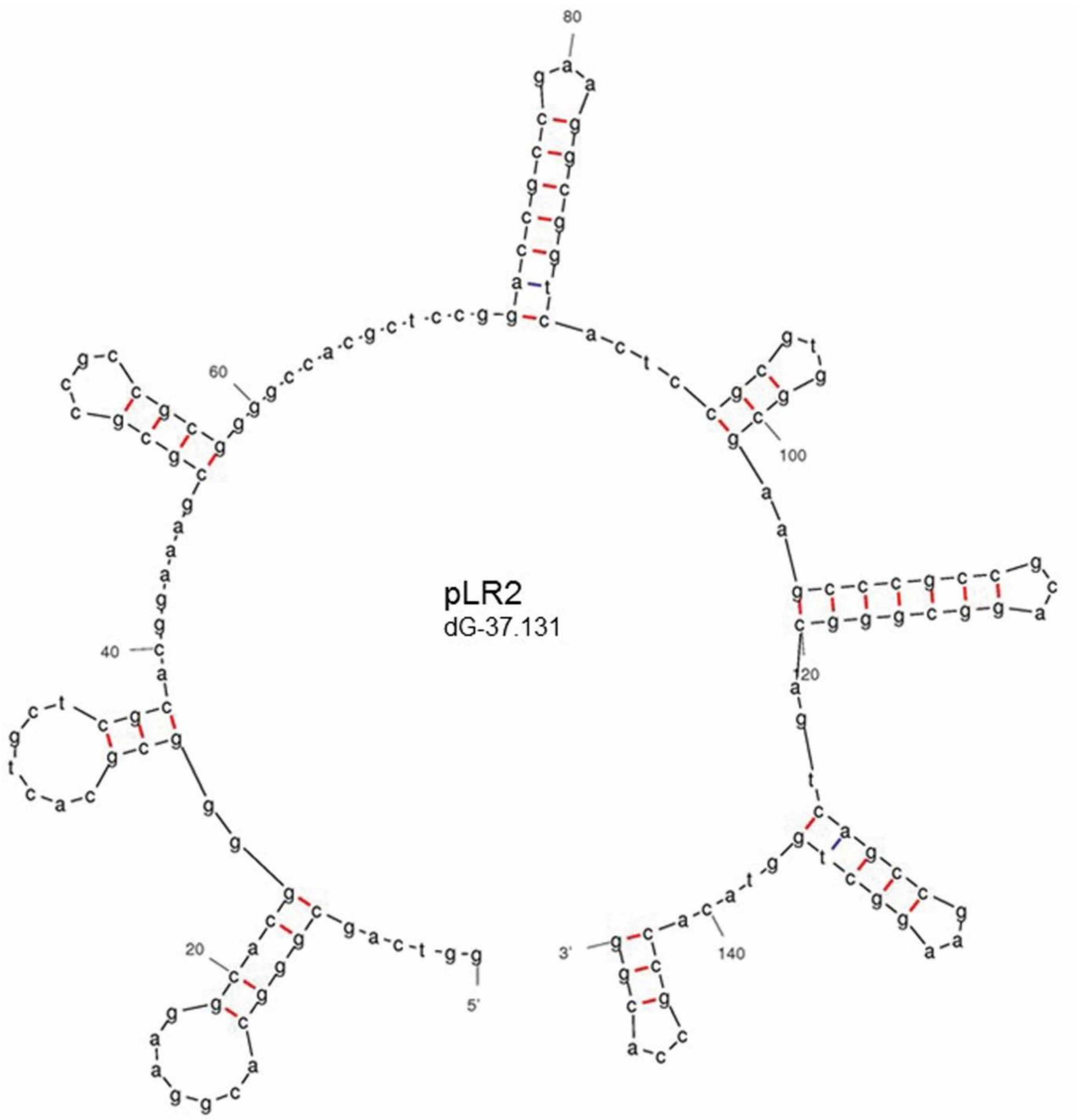


**Fig. S4. The six known classes of streptomycete telomeres**. Mfold projections of the terminal 150 bp of the known classes of streptomycete telomeres showing stem-loop structures (underlined, arrows) and hairpins (I-VIII, red) identified using Mfold (Zuker, 2003) are displayed where the revised free energies (ΔG) were determined using Jacobson-Stockmeyer theory to assign free energies to multi-branch loops. Projections were calculated using default conditions except folding temperature was set at 30°C, $Na^+$ concentration of 0.05M and maximum distance between paired bases was set at 20 (Yang et al., 2017). Sco, archetypal end from *S. coelicolor* chromosome (Bentley et al., 2002); SCP1, non-archetypal end from *S. coelicolor* plasmid SCP1 (Huang et al., 2007); Sg13350, chromosome end from

*S. griseus* 13350 chromosome (Ohnishi et al., 2008); Sg2247, chromosome end from *S. rimosus* ATCC10970 chromosome (this work); pRL1 end from *Streptomyces* sp. 44030 plasmid pRL1(Zhang et al., 2006); pRL2 end from *Streptomyce*s sp. 440414 plasmid pRL2 (Zhang et al., 2006).

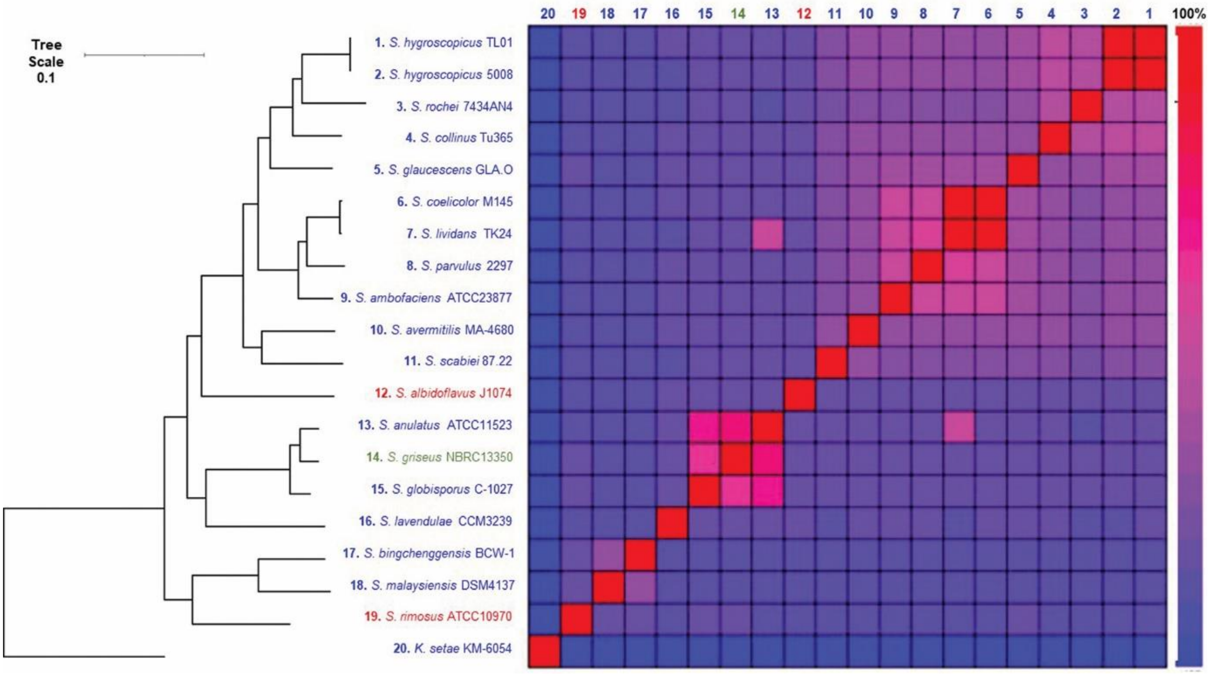


**Fig. S5. Phylogenetic relationship of the chromosomes from members of the *Streptomycetaceae* with closed genomes.**

Twenty closed streptomycete genomes (Table S2), where all replicons were completely sequenced and were either circular or, when linear, were flanked by one of the six classes of streptomycete telomeres, were arranged so that *dnaN* and *dnaA* were coded on the bottom stand and *parA* and *parB* on the top strands. The nineteen chromosome sequences from the genus *Streptomyces*, in conjunction with the closed sequence of *K. setae* KM-6054 as an outgroup, were then used to carry out Multi-Locus Sequence Analysis (MLSA) to produce a high-resolution species tree using AutoMLST after a concatenated alignment (Alanjary et al., 2019). All branches were supported with boot strap values of 100 except the *S. hygroscopicus*/*S. rochei* node (66) and *S. lividans*/*S. parvulus* node (98). Chromosomes with archetypal chromosomes are listed in blue, Sg2247 in red and Sg13350 in green. The average nucleotide identities (ANI) between the 20 closed *Streptomycetaceae* genomes was calculated using the OrthoANI tool (Lee et al., 2016) and plotted with the heat map function in R (Ver. 4.0).

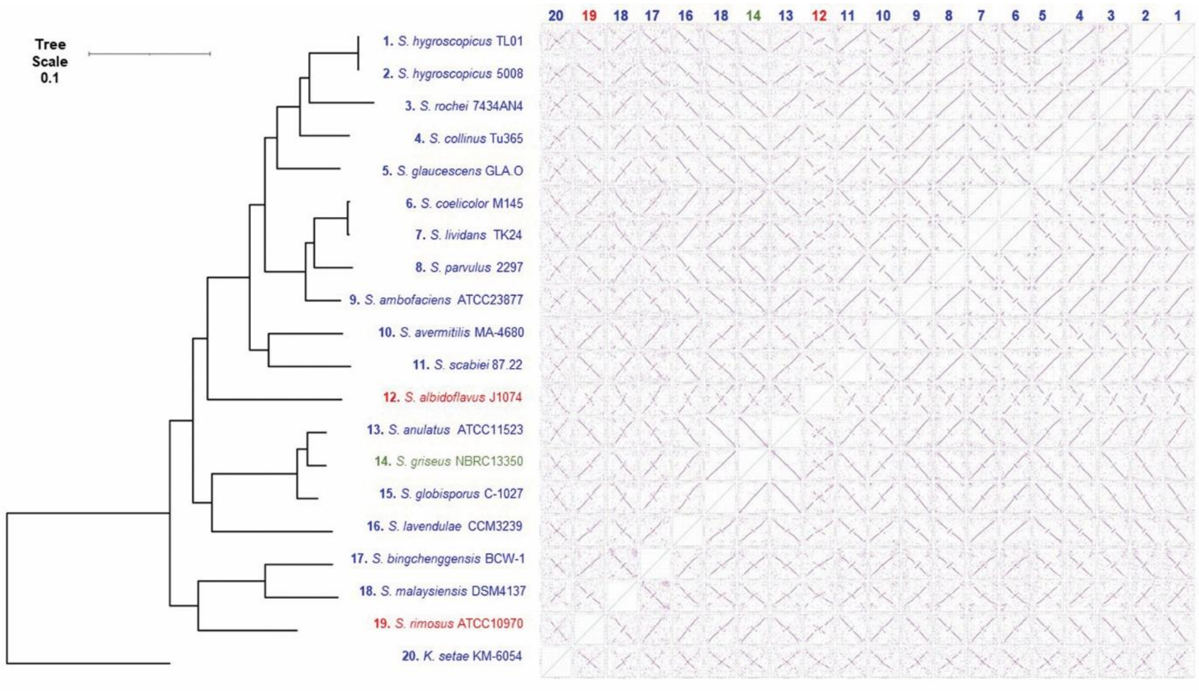


**Fig. S6. Symmetrical genomic rearrangements in streptomycete chromosomes.**

Dotplots comparing each of the 20 closed streptomycete chromosomes with each other, organized so that *oriC* regions were syntenous, were determined using Nucmer (Marcais et al., 2018). Reference sequences are displayed in columns (20-1) and query sequences in rows (1-20). Chromosomes with archetypal telomeres are listed in blue, Sg2247 telomeres in red and Sg13350 in green. Individual Nucmer plots of strains around selected nodes are displayed in more detail in Fig. 5.

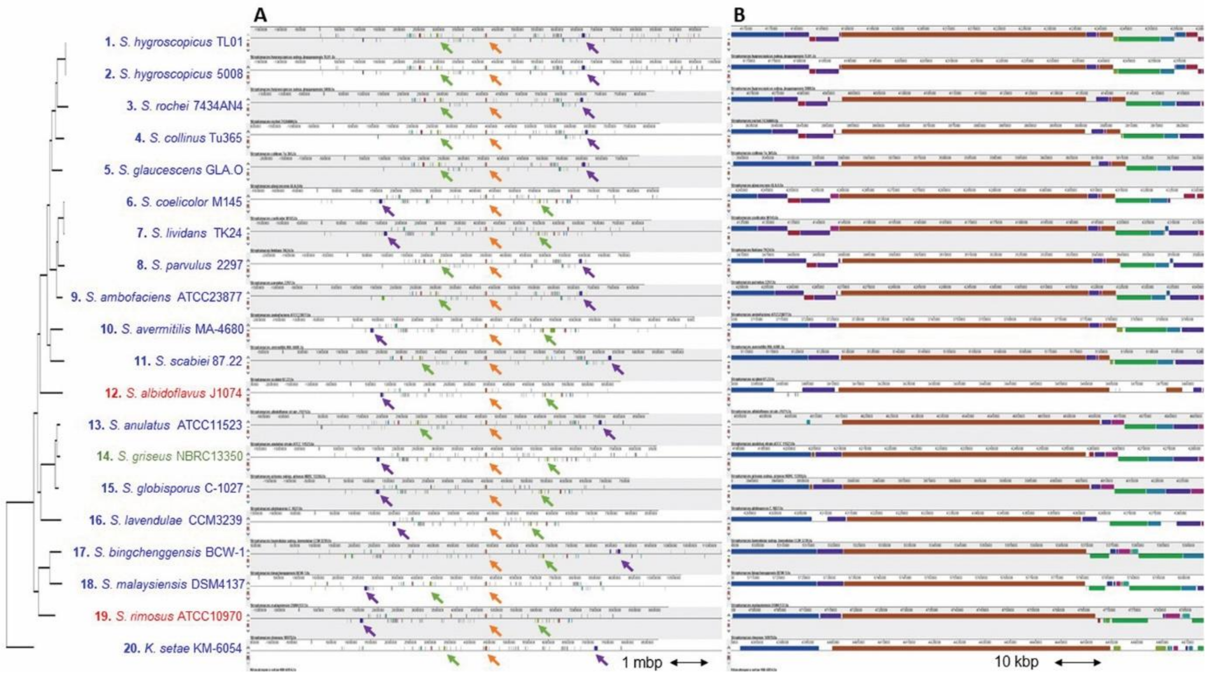


**Fig. S7. Streptomycete chromosomes are centred on a ~50kb origin island.**

All 20 closed streptomycete chromosomes were subjected to a progressive alignment in Mauve (Darling et al., 2010) using default settings. Replicons were centred on the *oriC* region of *S. coelicolor* M145 from *SCO3872-SCO3911* (brown locally collinear block (LCB) (brown arrow)) and displayed as the entire chromosome (A). Representative LCBs (green/purple LCBs/arrows) flanking the origin island showing their location in different replichores from different strains are also displayed. The *oriC* region is also displayed showing the conserved ~50-Kb origin island (B). This brown LCB corresponding to the *oriC* region of all chromosomes is flanked by a tRNA-ile (left) and *dnaB* (right, *SCO3911*) and represents the origin island. Chromosomes with archetypal telomeres are listed in blue, Sg2247 telomeres in red and Sg13350 in green.

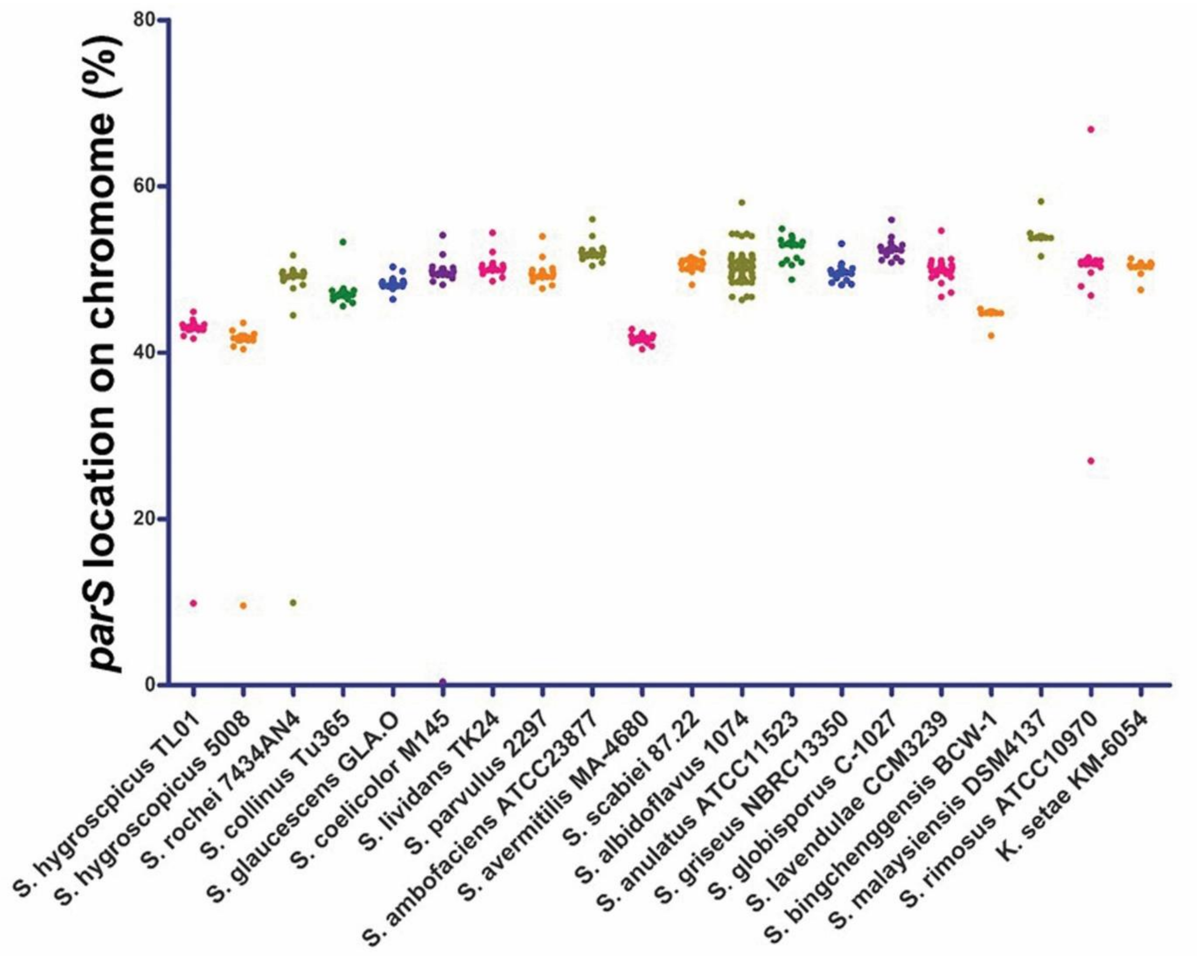


**Fig. S8. Location of *parS* sites on the closed genomes of the family *Streptomycetaceae* (all sites)**

The location of predicted *parS* sites were determined by searching the 20 closed genomes using the consensus matrix for bacterial *parS* sites (Livny et al., 2007). This was done by performing a ClustalW alignment of the 1030 predicted *parS* sited previously identified (Livny et al., 2007). A consensus matrix was then constructed in Weblogo (http://weblogo.threeplusone.com/create.cgi) (Crooks et al., 2004) and used to interrogate the 20 closedstreptomycete genomes using matrix scan in the Regulatory Sequence Analysis Tools suite (RSAT) (http://embnet.ccg.unam.mx/rsat//matrix-scan-quick_form.cgi) (Turatsinze et al., 2008). For this analysis the Threshold weight score was set at >15 (Livny et al., 2007). This allowed predicted *parS* sites to be mapped onto the chromosome and displayed as a scatter plot of the location of each *parS* site. Each location was normalised to genome size by expressing the site location as a percentage of the size of the chromosome.

**Table S1. iPCR primers for recovery of *S. rimosus* chromosomal and left & right plasmid telomeres.**

**Table S2 Known and predicted telomeres from the family *Streptomycetaceae***

Telomeres were identified on the basis of similarity with the six reported classes of streptomycete telomeres (archetypal (Sco), non-archetypal (SCP1), Sg2247, Sg13350, pLR1 and pLR2. The sequences are displayed in Fig. 1 were used as representatives of each telomeric class. The terminal 150bp or the sequence corresponding to the two terminal stem loop structures of each telomere class were used as query sequences in Blastn (NCBI). Hits were manually curated; those sequences that were either physically recovered and confirmed through Sanger sequencing or were found at the end of replicons were included as members of the four different telomeric classes. Telomeres were further sub-divided on the basis of their location (chromosome or linear plasmid; location at one or both ends of replicon) or means of identification (whole genome sequencing, physical recovery and traditional Sanger sequencing). Accession numbers for all sequences are displayed except that from *S. griseus* 2247 (not present in NCBI and only exists as a Figure in the article describing its recovery and characterisation (Goshi et al., 2002)).

**Table S3 Closed genomes from the family *Streptomycetatceae***

Organisms with closed genomes of the family *Streptomyetaceae* were defined as such on the basis that they are described as "complete" in NCBI and that all linear replicons found in that organism were flanked by known members of the six telomeric classes (Fig. 1) Archetypal (Sco), non-archetypal (SCP1), Sg13350, Sg2247, pLR1, pLR2. On this basis, although at time of writing 223 complete genomes from the family *Streptomycetaceae* are listed in NCBI, only 20 met our criteria as closed. Some organisms met some criteria for inclusion in our list of closed genomes, such as the chromosomes of *Streptomyces* sp. ETH9427 and *Streptomyces clavuligerus* F1D-5. However these organisms were not included in our closed list as these strains carried linear plasmids without recognizable telomeres. The length of the Terminal Inverted repeats at the ends of each replicon were determined by aligning each end of a replicon to the other end. This was done using the align sequence tool in Snapgene™ and allowed the identification of both perfect and imperfect repeat regions. Orthologues of genes from these 20 strains that encoded the proteins Tap (SCO7733, Tpg (SCO7734), TtrA (SCO0002), DnaN (SCO3878), DnaA (SCO3879), ParA (SCO3886), ParB (SCO3887) and RecA (SCO5769) were determined using BlastP (NCBI) with the proteins from *S . coelicolor* A3(2) as queries; only hits with at least 25% identity and coverage were included.

**Table S4 Origin islands and *parS* sites from the family *Streptomycetaceae***

The location of predicted *parS* sites with respect to origin islands on the chromosomes of the 20 closed genomes from the family *Streptomycetaceae* were determined by searching the 20 these genomes using the consensus matrix for bacterial *parS* sites (Livny et al., 2007). This was done by performing a ClustalW alignment of the 1030 predicted *parS* sited previously identified (Livny et al., 2007). A consensus matrix was then constructed in Weblogo (http://weblogo.threeplusone.com/create.cgi) (Crooks et al., 2004) and used to interrogate the 20 closed *Streptomyetaceae* genomes using matrix scan in the Regulatory Sequence Analysis Tools suite (RSAT) (http://embnet.ccg.unam.mx/rsat//matrix-scan-quick_form.cgi) (Turatsinze et al., 2008). For this analysis the Threshold weight score was set at >15 (Livny et al., 2007). This allowed predicted *parS* sites to be mapped onto the chromosome and their locations determined with respect to the origin island and in relation to *oriC*. Each location was also normalised to genome size by expressing the site location as a percentage of the size of the chromosome. In *S. coelicolor* the origin island contains the following genes. tRNA-ile, *SCO3872, SCO3873* (*gyrA*)*, SCO3874* (*gyrB*)*, SCO3875, SCO3876* (*recF*)*, SCO3877, SCO3878* (*dnaN*)*, oriC, SCO3879* (*dnaA*)*, SCO3880* (*rpmH*)*, SCO3881* (*rnpA*)*, SCO3882, SCO3883* (*yidC*)*, SCO3884, SCO3885* (*gidB*)*, SCO3886* (*parA*)*, SCO3887* (*parB*)*, SCO3888, SCO3889* (*trxA*)*, SCO3890* (*trxB*)*, SCO3891, SCO3892* (*sigT*)*, SCO3893, SCO3894, SCO3895, SCO3896, SCO3897, SCO3898, SCO3899, SCO3900, SCO3901, SCO3902, SCO3903, SCO3904, SCO3905, SCO3906* (*rpsF*)*, SCO3907 (ssbA), SCO3908* (*rpsR*)*, SCO3909 (rplI), SCO3910, SCO3911 (dnaB).* In *S. coelicolor oriC* lies between *SCO3878* (*dnaN*) and *SCO3879* (*dnaA*), after arranging all chromosomes so that these genes were transcribed in right to left direction, the last base of *dnaA* from all genomes was selected as the location of *oriC*.


ALANJARY, M., STEINKE, K. & ZIEMERT, N. 2019. AutoMLST: an automated web server for generating multi-locus species trees highlighting natural product potential. *Nucleic Acids Res,* 47**,** W276-W282.

BENTLEY, S. D., CHATER, K. F., CERDENO-TARRAGA, A. M., CHALLIS, G. L., THOMSON, N. R. & JAMES, K. D. 2002. Complete genome sequence of the model actinomycete Streptomyces coelicolor A3(2). *Nature,* 417.

CROOKS, G. E., HON, G., CHANDONIA, J.-M. & BRENNER, S. E. 2004. WebLogo: A Sequence Logo Generator. *Genome Research,* 14**,** 1188-1190.

DARLING, A. E., MAU, B. & PERNA, N. T. 2010. progressiveMauve: multiple genome alignment with gene gain, loss and rearrangement. *PLoS One,* 5**,** e11147.

FAN, Y., DAI, Y., CHENG, Q., ZHANG, G., ZHANG, D., FANG, P., WU, H., BAI, L., DENG, Z. & QIN, Z. 2012. A self-ligation method for PCR-sequencing the telomeres of Streptomyces and Mycobacterium linear replicons. *J Microbiol Methods,* 90**,** 105-7.

GOSHI, K., UCHIDA, T., LEZHAVA, A., YAMASAKI, M., HIRATSU, K., SHINKAWA, H. & KINASHI, H. 2002. Cloning and analysis of the telomere and terminal inverted repeat of the linear chromosome of Streptomyces griseus. *J Bacteriol,* 184**,** 3411-5.

HUANG, C. H., TSAI, H. H., TSAY, Y. G., CHIEN, Y. N., WANG, S. L., CHENG, M. Y., KE, C. H. & CHEN, C. W. 2007. The telomere system of the Streptomyces linear plasmid SCP1 represents a novel class. *Mol Microbiol,* 63**,** 1710-8.

LEE, I., OUK KIM, Y., PARK, S. C. & CHUN, J. 2016. OrthoANI: An improved algorithm and software for calculating average nucleotide identity. *Int J Syst Evol Microbiol,* 66**,** 1100-1103.

LIVNY, J., YAMAICHI, Y. & WALDOR, M. K. 2007. Distribution of centromere-like parS sites in bacteria: insights from comparative genomics. *J Bacteriol,* 189**,** 8693-703.

MARCAIS, G., DELCHER, A. L., PHILLIPPY, A. M., COSTON, R., SALZBERG, S. L. & ZIMIN, A. 2018. MUMmer4: A fast and versatile genome alignment system. *PLoS Comput Biol,* 14**,** e1005944.

OHNISHI, Y., ISHIKAWA, J., HARA, H., SUZUKI, H., IKENOYA, M., IKEDA, H., YAMASHITA, A., HATTORI, M. & HORINOUCHI, S. 2008. Genome sequence of the streptomycin-producing microorganism Streptomyces griseus IFO 13350. *J Bacteriol,* 190**,** 4050-60.

TURATSINZE, J.-V., THOMAS-CHOLLIER, M., DEFRANCE, M. & VAN HELDEN, J. 2008. Using RSAT to scan genome sequences for transcription factor binding sites and cis-regulatory modules. *Nature Protocols,* 3**,** 1578-1588.

YANG, C. C., TSENG, S. M., PAN, H. Y., HUANG, C. H. & CHEN, C. W. 2017. Telomere associated primase Tap repairs truncated telomeres of Streptomyces. *Nucleic Acids Res,* 45**,** 5838-5849.

ZHANG, R., YANG, Y., FANG, P., JIANG, C., XU, L., ZHU, Y., SHEN, M., XIA, H., ZHAO, J., CHEN, T. & QIN, Z. 2006. Diversity of telomere palindromic sequences and replication genes among Streptomyces linear plasmids. *Appl Environ Microbiol,* 72**,** 5728-33.

ZUKER, M. 2003. Mfold web server for nucleic acid folding and hybridization prediction. *Nucleic Acids Res,* 31**,** 3406-15.
