## Supplementary Figs and Files for "Bilateral symmetry of linear streptomycete chromosomes": Bilateral Xsomes Supp Info Final.pdf

<u>**Supplementary Information**</u>

### Assembly of the *S. rimosus* ATCC 10970 closed genome sequence

The complete genome sequence of *S. rimosus* ATCC10970, was assembled using a combination of reads from long-read PacBio sequencing and short-read Illumina sequencing. In the first instance, assembly of PacBio data was carried out through HGAP using CCS reads generated from this raw data, with a minimum of 3 reads through a single molecule; this yielded 19555 sequences between 48 and 16693bp in length. Meanwhile Illumina sequencing generated 2,415,879 reads with a mean coverage of 101.793x.

PacBio sequences were assembled into five contigs using HGAP (Contig A, 3,195,161bp; Contig B, 2,836,700bp; Contig C, 2,083,116bp; Contig D, 1, 240,339bp and Contig E, 278,157bp). Contig E was identified as the known *S. rimosus* G7 Giant Linear Plasmid (GLP) due to its similarity in size to the known GLP of this strain; estimated to be slightly smaller than the 312Kb GLP of *S. rimosus* R6 determined by pulsed-field gel electrophoresis (PFGE) (Gravius et al., 1994). In addition, this contig also contained copies of *parA*, *parB* and *traSA* characteristic of streptomycete GLPs (Thoma et al., 2016).

Contigs A-D were joined using data from publically available data from two *S. rimosus* draft genome sequences (*S. rimosus* ATCC 10970, DDBJ/EMBL/GenBank GenBank accession number: ANSJ00000000 (Pethick et al., 2013) and *S. rimosus* NRRL ISP5260, DDBJ/EMBL/GenBank: JNYR00000000.1). Contigs A and B were joined using the bridging contig NZ_ANSJ01000266.1 and Contigs B and C using the bridging contig NZ_ANSJ01000079.1|. The other end of Contig C carried an rRNA operon as did the telomere-distal end of Contig D, but we were unable to identify existing overlapping contigs from other sequencing projects. As a result, the hybrid assembler Unicycler was used to generate an assembly of both long PacBio and short Illumina sequences and generated 56 contigs; one of these contigs (Contig 2, 1,201,767bp) spanned Contigs C and D allowing assembly the chromosome as a single contig. Contig A and Contig D contained ~11Kb Terminal Inverted Repeat (TIRs) which suggested that these contigs represented the terminal contigs. This was confirmed by the identification of two contigs, NZ_JNYR01000065 and NZ_JNYR01000071, that overlapped with the TIRs of Contig A and Contig D respectively. This allowed us to further extend the sequence of the TIRs. The sequences of both of these two contigs were also found at the *oriC*-distal ends of both Contig C and Contig D. Contig E was extended through identification of the bridging contigs NZ_ANSJ01000106.1| and NZ_ANSJ01000002.1. Despite this, the ends of the draft linear chromosome did not contain either archetypal or non-archetypal telomeres. As a result we independently recovered and sequenced the telomeres of both the GLP and chromosome (see materials and methods). Comparison of the telomeric sequences with our assembled contiguous chromosomal and plasmid contigs allowed us to extend the contiguous sequences to close the sequence of both replicons.

Following the construction of a draft sequence, the sequence was polished by first using Bowtie (Galaxy version 1.2.0) (Langmead et al., 2009) where Illumina paired end reads (FastQ files were aligned with the draft sequence to generate a BAM files. This file was then mapped to the draft sequence files (both chromosome and plasmid) using Pilon in Galaxy Version 1.20.1,(Walker et al., 2014) to generate the final sequences of the two replicons. Annotation was carried out using NCBI Prokaryotic Genome Annotation Pipeline (PGAP) and is available as Bioproject PRJNA182749 and Biosample SAMN02471950. The *S. rimosus* ATCC 10970 chromosome is listed under accession number CP048261 and the plasmid, pSRP1, CP048262.


GRAVIUS, B., GLOCKER, D., PIGAC, J., PANDŽA, K., HRANUELI, D. & CULLUM, J. 1994. The 387 kb linear plasmid pPZG101 of *Streptomyces rimosus* and its interactions with the chromosome. *Microbiology,* 140**,** 2271-2277.

LANGMEAD, B., TRAPNELL, C., POP, M. & SALZBERG, S. L. 2009. Ultrafast and memory-efficient alignment of short DNA sequences to the human genome. *Genome Biology,* 10**,** R25.

PETHICK, F. E., MACFADYEN, A. C., TANG, Z., SANGAL, V., LIU, T. T., CHU, J., KOSEC, G., PETKOVIC, H., GUO, M., KIRBY, R., HOSKISSON, P. A., HERRON, P. R. & HUNTER, I. S. 2013. Draft Genome Sequence of the Oxytetracycline-Producing Bacterium *Streptomyces rimosus* ATCC 10970. *Genome Announc,* 1**,** e0006313.

THOMA, L., VOLLMER, B. & MUTH, G. 2016. Fluorescence microscopy of *Streptomyces* conjugation suggests DNA-transfer at the lateral walls and reveals the spreading of the plasmid in the recipient mycelium. *Environ Microbiol,* 18**,** 598-608.

WALKER, B. J., ABEEL, T., SHEA, T., PRIEST, M., ABOUELLIEL, A., SAKTHIKUMAR, S., CUOMO, C. A., ZENG, Q., WORTMAN, J., YOUNG, S. K. & EARL, A. M. 2014. Pilon: an integrated tool for comprehensive microbial variant detection and genome assembly improvement. *PLoS One,* 9**,** e112963.
