## Supplementary Figs and Files for "Bilateral symmetry of linear streptomycete chromosomes": Table S1. iPCR primers for recovery of S. rimosus chromosomal and left & right plasmid telomeres.pdf

**Table S1. iPCR primers for recovery of *S. rimosus* chromosomal (Lis-SrimC-iPCR-1 & 2) and left (Lis-SrimP-iPCR-L1 & L2) & right plasmid telomeres (Lis-SrimP-iPCR-R1 & L2)**

| Name | Sequence (5'-3') |
|---|---|
| Lis-SrimC-iPCR-1 | ttgcaaaaatcgtcggttgggagtcggt |
| Lis-SrimC-iPCR-2 | catcacgctgcatcagttttggagaaacgt |
| Lis-SrimP-iPCR-L1 | atgacgcgattttccgaggcctttccgta |
| Lis-SrimP-iPCR-L2 | ccggagggtttattatctttgcggcccgta |
| Lis-SrimP-iPCR-R1 | gccttgcatcatgctgcatcagtttcgggaaaa |
| Lis-SrimP-iPCR-R2 | agaacaatgcgggaaagtcgagcgtgatc |
